# Ammonia-oxidizing archaea possess a wide range of cellular ammonia affinities

**DOI:** 10.1101/2021.03.02.433310

**Authors:** Man-Young Jung, Christopher J. Sedlacek, K. Dimitri Kits, Anna J. Mueller, Sung-Keun Rhee, Linda Hink, Graeme W. Nicol, Barbara Bayer, Laura Lehtovirta-Morley, Chloe Wright, Jose R. de la Torre, Craig W. Herbold, Petra Pjevac, Holger Daims, Michael Wagner

## Abstract

Nitrification, the oxidation of ammonia to nitrate, is an essential process in the biogeochemical nitrogen cycle. The first step of nitrification, ammonia oxidation, is performed by three, often co-occurring guilds of chemolithoautotrophs: ammonia-oxidizing bacteria (AOB), archaea (AOA), and complete ammonia oxidizers (comammox). Substrate kinetics are considered to be a major niche-differentiating factor between these guilds, but few AOA strains have been kinetically characterized. Here, the ammonia oxidation kinetic properties of 12 AOA representing all major phylogenetic lineages were determined using microrespirometry. Members of the genus *Nitrosocosmicus* have the lowest substrate affinity of any characterized AOA, which are similar to previously determined affinities of AOB. This contrasts previous assumptions that all AOA possess much higher substrate affinities than their comammox or AOB counterparts. The substrate affinity of ammonia oxidizers correlated with their cell surface area to volume ratios. In addition, kinetic measurements across a range of pH values strongly supports the hypothesis that – like for AOB – ammonia and not ammonium is the substrate for the ammonia monooxygenase enzyme of AOA and comammox. Together, these data will facilitate predictions and interpretation of ammonia oxidizer community structures and provide a robust basis for establishing testable hypotheses on competition between AOB, AOA, and comammox.

## Introduction

Nitrification, the microbially mediated oxidation of ammonia (NH3) to nitrate (NO3^-^) via nitrite (NO2^-^), is a key process of the biogeochemical nitrogen cycle ^1, 2^ and is mostly driven by autotrophic microorganisms that are capable of growing with NH3 and/or NO2^-^ as sole energy generating substrates. For more than a century, ammonia-oxidizing bacteria (AOB) were considered the lone drivers of aerobic ammonia oxidation by autotrophs, as ammonia-oxidizing archaea (AOA) ^3, 4^ and complete ammonia oxidizers (comammox) ^5–7^ eluded discovery until relatively recently. Our present-day understanding of ammonia oxidation is quite different: AOA frequently outnumber AOB in oligotrophic habitats ^8–10^, while AOB often dominate in eutrophic environments ^11–14^. Comammox have been shown to be abundant and even dominant in various natural and engineered environments ^15–19^, although the habitat range and ecophysiology of comammox remains less well resolved. Notably, in the majority of ecosystems – with the exception of the marine environment, where no comammox has been detected – AOA, AOB, and comammox often co-occur.

Many environmental and physiological factors are known to affect the niche differentiation and habitat selection of ammonia-oxidizing microorganisms (AOM) ^20, 21^. In fact, AOM species display differential responses to factors such as pH, oxygen concentrations, light conditions, temperature, metal and organic compounds, and substrate concentrations ^22–27^. These differential responses are frequently used to explain the co-occurence of AOM across environments. However, the cellular properties underlying these niche-differentiating physiological characterstics of AOM often remain unclear.

The substrate affinity of a microorganism can be expressed with Michaelis-Menten kinetic equations, analogous to enzyme kinetics, defined by an apparent-half-saturation activity (*K*_m(app))_ and a maximal activity rate (*V*_max_). In addition, the specific substrate affinity (*a*_o_) takes into account both the cellular *K*_m(app)_ and *V*_max_, and is thus an appropriate measure for comparing interspecies competitiveness _28_. Based on whole cell kinetic properties, AOM were observed to have different survival or lifestyle strategies. The first study investigating the whole cell kinetics of an AOA revealed that *Nitrosopumilus maritimus* SCM1 displayed a low maximum NH_3_ oxidation activity rate, but a very high substrate affinity for NH3 and *a*°, compared with AOB ^29^. Based on these findings with a single AOA strain, substrate affinity was postulated as a major niche-differentiating factor between AOA and AOB ^20, 29^. However, recently it was shown that (i) the only comammox isolate *Nitrospira inopinata* has a *K*_m(app)_ for NH_3_ lower than that of all characterized AOB and (ii) that the *K*_m(app)_ for NH_3_ in a few non-marine AOA strains is not always orders of magnitude lower than that of AOB_5_. Nevertheless, the AOA with comparatively high *K*_m(app)_ for NH_3_ (low affinity) still possess a significantly higher *a*° than AOB, indicating that these AOA are still more efficient substrate scavengers^5^. Furthermore, temperature and pH, which are known niche-differentiating factors ^30–32^, have previously been shown to affect the substrate affinity of AOB ^33–35^, but the influence of these parameters on the substrate affinity of AOA and comammox remains to be determined.

In this study, the whole cell kinetic properties of twelve AOA species were determined through instantaneous substrate-dependent microrespirometry (MR) experiments. These include representatives from all four major AOA phylogenetic lineages, isolated or enriched from various habitats (i.e. marine, terrestrial, and geothermal) and possessing a wide variety of pH and temperature growth optima. In these analyses, we also explored the links between the cellular *K*_m(app)_ and *a*° of AOM with their cell surface area to volume ratio. Furthermore, by performing riments at different pH values we investigated whether the undissociated NH3 or m (NH4+) is the substrate for AOA and comammox.

## Results and Discussion

### AOA kinetic properties

In this study we investigated the kinetic properties of twelve AOA strains, including representatives from all four described AOA phylogenetic lineages: *Nitrosopumilales* (Group I.1a), ‘*Ca.* Nitrosotaleales’ (Group I.1a-associated), *Nitrososphaerales* (Group I.1b), and ‘*Ca.* Nitrosocaldales’ (thermophilic AOA clade) ^36, 37^ (Fig. 1). These AOA isolates and enrichments were obtained from a variety of habitats (marine, soil, sediment, hot spring) and have optimal growth pH and temperatures ranging from 5.3-7.8 and 25-72°C, respectively (Supplementary Table 2). The substrate-dependent oxygen consumption rates for all AOA tested followed Michaelis-Menten kinetics. Below, the kinetic properties of these AOA are put into a broader context with comparisons to previously characterized AOM.

**Fig. 1.**
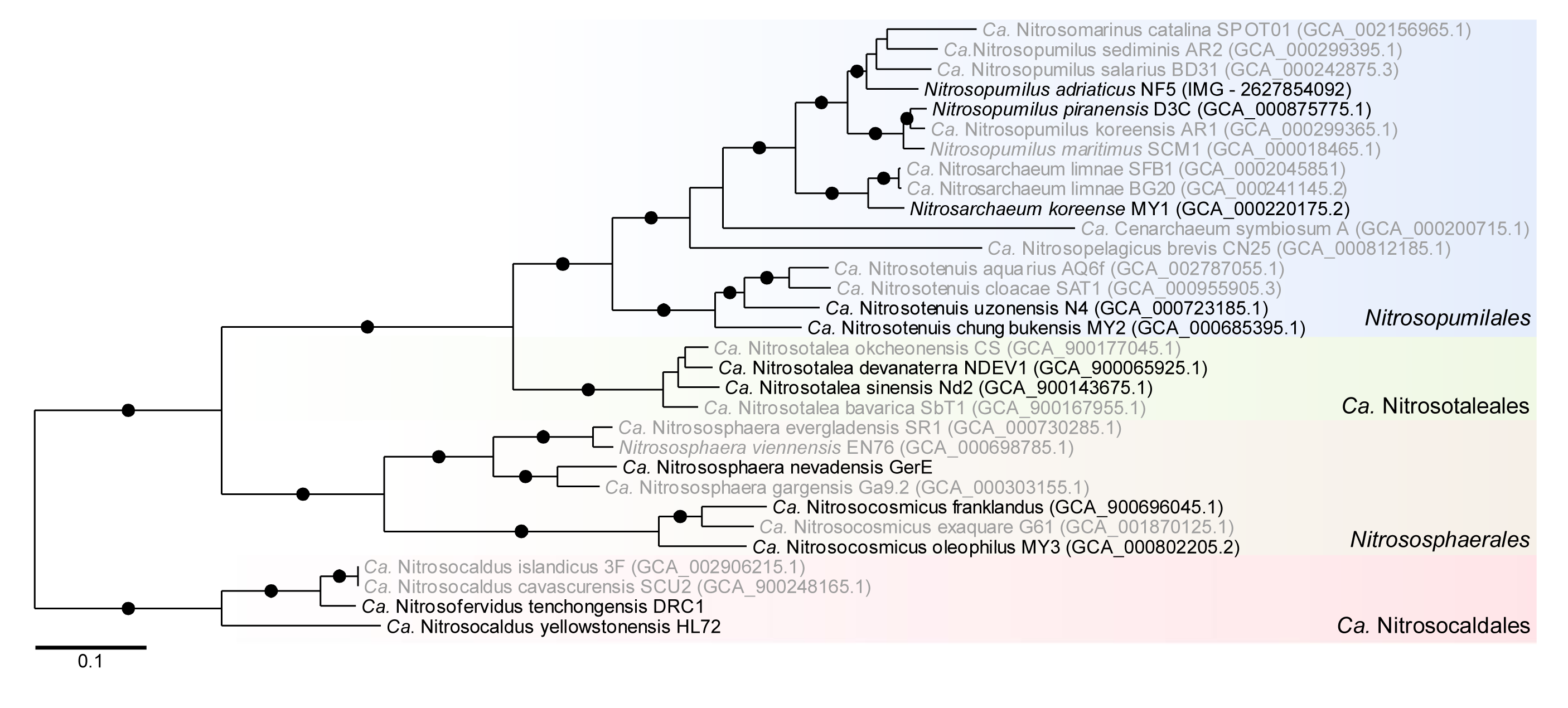
A phylogenetic reconstruction of the AOA used in this study. AOA kinetically characterized for the first time are shown in black. ‘*Ca*. N. uzonensis’ N4 is also labeled in black as its previous kinetic characterization occurred as an enrichment culture ^5^. An unrooted approximate maximum likelihood tree was calculated using IQ-TREE under model LG+F+R5 using an alignment of 34 universal genes (43 markers). Support values (UFboot) greater than 95% for bipartitions are shown with a filled circle. All other bipartitions received <80% UFboot support. The tree was arbitrarily rooted on the branch leading to the *Nitrosocaldaceae* and order designations reflect lineages proposed by Alves *et al*., 2018. The scale bar indicates amino acids changes per site.

### Nitrosopumilales (Group I.1a)

From this lineage, three mesophilic marine (*N. piranensis* D3C, *N. adriaticus* NF5, and *N. maritimus* SCM1) _3,38_, two agricultural soil (*N. koreense* MY1 and ‘*Ca.* N. chungbukensis’ MY2) _39,40_ and one thermal spring isolate (‘*Ca.* N. uzonensis’ N4)^41^ were kinetically characterized (Supplementary Fig. 1). These AOA all displayed a high substrate affinity for NH3, ranging from ∼2.2 to 24.8 nM. Thus, all characterized *Nitrosopumilales*, and not just marine isolates, are adapted to oligotrophic conditions. All possess substrate affinities several orders of magnitude higher (lower *K*m(app)) than any characterized AOB, with the exception of the recently characterized acidophilic gammaproteobacterial AOB ‘*Ca.* Nitrosacidococcus tergens’ ^42^ (Fig. 2a). This finding appears to support the widely reported hypothesis that regardless of the environment, AOA in general are adapted to lower substrate concentrations than AOB ^22, 29, 30^. However, as described later, this trend does not apply to all AOA.

**Fig. 2.**
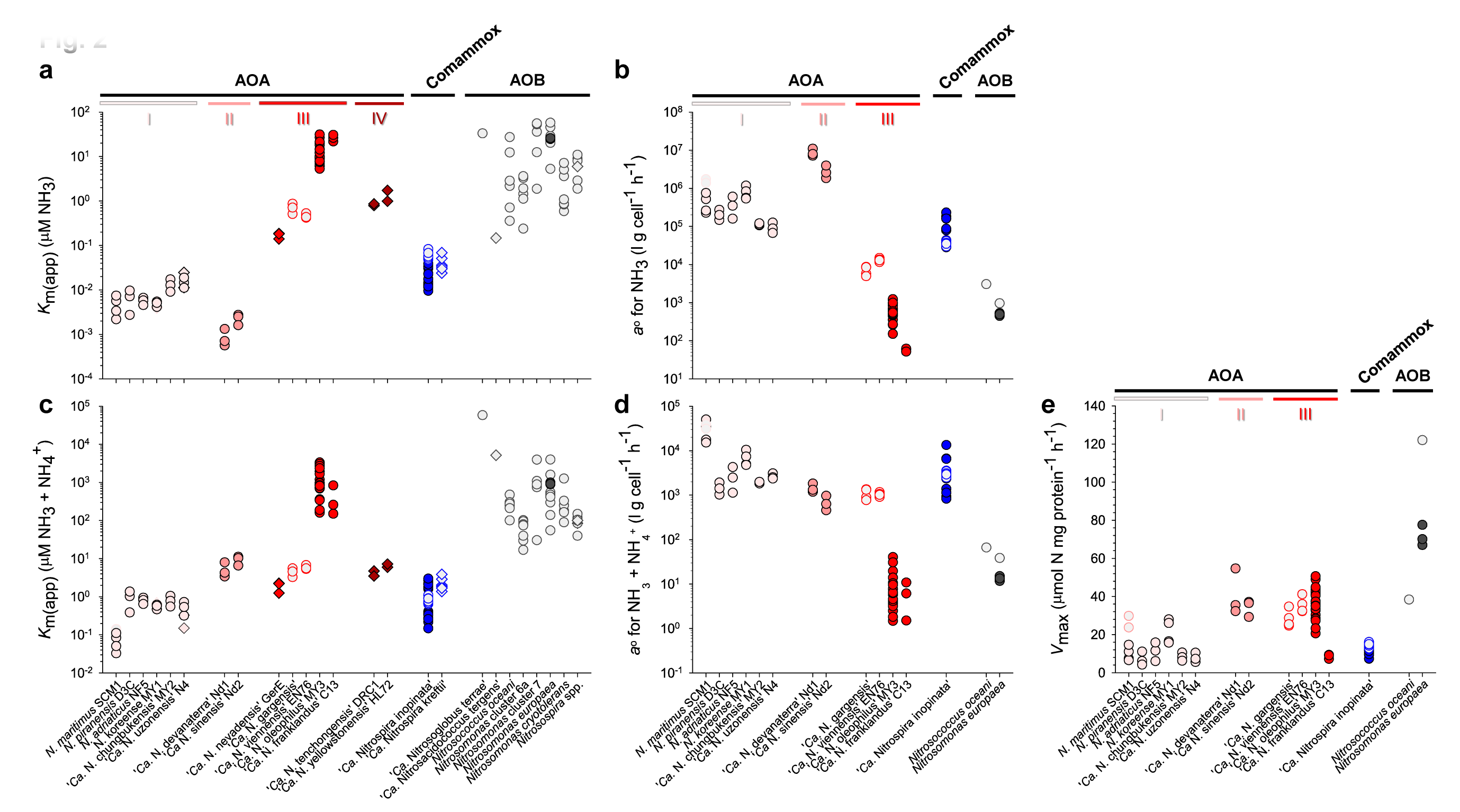
Substrate-dependent oxidation kinetics of ammonia-oxidizing microorganisms. The (a) apparent substrate affinity (*K*_m(app))_ for NH3, (b) specific substrate affinity (*a*°) for NH3, (c) *K*m(app) for total ammonium, (d) *a*° for total ammonium, and (e) maximum oxidation rate (*V*max), of AOA (red), comammox (blue), and AOB (black) are provided. Symbols filled with light grey represent previously published values from reference studies (references provided in Materials and Methods). The four different gradations of red differentiate the four AOA phylogenetic lineages: (I) *Nitrosopumilales*, (II) ‘*Ca*. Nitrosotaleales’, (III) *Nitrososphaerales*, and (IV) ‘*Ca*. Nitrosocaldales’. Measurements were performed with either pure (circles) or enrichment (diamonds) cultures. Multiple symbols per strain represent independent measurements performed in this study and/or in the literature. The individual Michaelis-Menten plots for each AOM determined in this study are presented in Supplementary Figs. 1, 3-5, and 8. Note the different scales.

As the substrate oxidation kinetics of the marine AOA strain, *N. maritimus* SCM1, originally characterized by Martens-Habbena *et al*. ^29^ have recently been disputed ^43^, they were revisited in this study (Supplementary Fig. 2). With the same strain of *N. maritimus* used in Hink *et al*., ^43^ (directly obtained by the authors), we were able to reproduce (Supplementary Figs. 1 and 1. 2) the original kinetic properties of *N. maritimus* SCM1 reported in Martens-Habbena *et al.* ^29^ ruling out strain domestication during lab propagation as cause for the observed discrepancy. Therefore, the reported differences in the literature possibly reflect the measurements of two distinct cellular properties, *K*m(app) ^29^ and *K*s ^43^, representing the half saturation of activity and growth, respectively. In addition, differences in pre-measurement cultivation and growth conditions could also contribute to these unexpected differences ^43, 44^. More details are provided in the Supplementary Results and Discussion.

### ‘*Ca.* Nitrosotaleales’ (Group I.1a-associated)

The only isolated AOA strains in this lineage ‘*Ca.* Nitrosotalea devanaterra’ Nd1 and ‘*Ca.* Nitrosotalea sinensis’ Nd2, are highly adapted for survival in acidic environments and grow optimally at pH 5.3 ^25, 45^. Both display a relatively low affinity for total ammonium (*K*m(app) = 3.41 to 11.23 μM), but their affinity for NH3 is among the highest calculated of any AOA characterized (*K*m(app) = ∼0.6 to 2.8 nM) (Fig. 2a,c, and Supplementary Fig. 3). This seemingly drastic difference in substrate affinity for total ammonium versus NH3 is due to the high acid dissociation constant of ammonium (p*K*a = 9.25). The very limited availability of NH3 under acidic conditions has led to the hypothesis that these acidophilic AOA should be highly adapted to very low NH3 concentrations and possess a high substrate affinity (low *K*m(app)) for NH3 ^46, 47^. Our data corroborate this hypothesis.

### Nitrososphaerales (Group I.1b)

The AOA strains ‘*Ca.* N. nevadensis’ GerE (culture information provided in Supplementary Results and Discussion), ‘*Ca.* N. oleophilus’ MY3 ^48^ and ‘*Ca.* N. franklandus’ C13 ^49^ were kinetically characterized, and contextualized with the previously published kinetic characterization of *Nitrososphaera viennensis* EN76 and ‘*Ca.* Nitrososphaera gargensis’ ^5^. Together, the *Nitrososphaerales* AOA possess a wide range of affinities for NH3 (*K*m(app) = ∼0.14 to 31.5 µ M) (Fig. 2a and Supplementary Fig. 4). Although this range of NH3 affinities spans more than two orders of magnitude, none of the *Nitrososphaerales* AOA possess an affinity for NH3 as high as any *Nitrosopumilales* or *‘Ca.* Nitrosotaleales*’* AOA (Fig. 2a).

The moderately thermophilic enrichment culture ‘*Ca.* N. nevadensis’ GerE displayed a higher substrate affinity (lower *K*m(app)) for NH3 (0.17 ± 0.03 µ M) than the other characterized AOA strains within the genus *Nitrososphaera* (Fig. 2a). In contrast, ‘*Ca.* N. oleophilus’ MY3 and ‘*Ca.* N. franklandus’ C13, which belong to the genus *Nitrosocosmicus*, had the lowest affinity (highest *K*m(app)) for NH3 (12.37 ± 6.78 μM and 16.32 ± 14.11 μM, respectively) of any AOA characterized to date. In fact, their substrate affinity is comparable to several characterized AOB (Fig. 2a). In this context it is interesting to note that several *Nitrosocosmicus* species have been shown to tolerate very high ammonium concentrations ^48–50^, a trait usually associated with AOB ^24, 51^. The low substrate affinity observed in *Nitrosocosmicus* AOA correlates with the absence of a putative Amt-type high affinity ammonium transporter in the genome of any sequenced *Nitrosocosmicus* species to date ^48, 49, 52^.

### ‘Ca. Nitrosocaldales’ (Thermophilic AOA lineage)

The thermophilic AOA enrichment cultures, ‘*Ca.* Nitrosocaldus yellowstonensis’ HL72 ^53^ and ‘*Ca.* N. tenchongensis’ DRC1 (culture information provided in Supplementary Results and Discussion), possess affinities for NH3 (*K*m(app) = ∼1.36 ± 0.53 μM and ∼0.83 ± 0.01 μM) for NH3 comparable to AOA within the genus *Nitrososphaera* (Fig. 2a). Notably, the substrate oxidation rate of these two AOA quickly dropped with increasing substrate concentrations after *V*max was reached (Supplementary Fig. 5). This trend was not observed with any other AOA tested here and may reflect an increased susceptibly to NH3 stress at high temperatures, as the free NH3 concentration increases with increasing temperatures^33^.

Together, these results highlight that the substrate affinity for NH3 among AOA species is much more variable than previously hypothesized, spanning several orders of magnitude and in some cases overlapping with the substrate affinity values of characterized non-oligotrophic AOB. In addition, the substrate affinity of AOA is related, to a certain degree, to their phylogenetic placement within each of the four AOA phylogenetic lineages mentioned above (Fig. 2). Although the substrate affinity ranges of these AOA lineages overlap, the link between AOA phylogeny and kinetic properties provides deeper insights into the physiological and evolutionary differences among AOA species. While substrate affinity is certainly one of multiple factors that contribute to niche differentiation between AOM, it is more complex than the prevailing dogma that all AOA have a higher substrate affinity than their AOB counterparts and may be as significant in differentiating the niches of different AOA species as between AOA and AOB.

### Maximum substrate oxidation rates (*V*max)

The normalized maximum substrate oxidation rate of all the AOA characterized to date only span about one order of magnitude from 4.27 to 54.68 μmol N mg protein^-1^ h^-1^. These normalized AOA *V*max values are in the same range as the recorded *V*max for the comammox *N. inopinata* (∼12 μmol N mg protein^-1^ h^-1^) and the marine AOB strain *Nitrosococcus oceani* ATCC 19707 (∼38 μmol N mg protein^-1^ h^-1^) but are lower than the normalized *V*max of the AOB *Nitrosomonas europaea* ATCC 19718 (average of 84.2 μmol N mg protein^-1^ h^-1^; Fig. 2e). The high *V*max value for *N. europaea* is the only real outlier among the AOM characterized to date and it remains to be determined whether other AOB related to *N. europaea* also possess such a high *V*max or if members of the *Nitrosomonadales* possess a broad range of *V*max values. Similarly, as additional comammox strains become available as pure cultures their kinetic characterization will be vital in understanding the variability of these ecologically important parameters within this guild.

### Specific substrate affinity (*a*°)

Although the *K*m(app) and *V*max of AOM can be compared by themselves and provide useful information on cellular properties, the ability of an AOM to scavenge (and compete for) substrate from a dilute solution is most appropriately represented by the *a*°, which takes into account both the cellular *K*m(app) and *V*max ^28^. In previous studies, the *a*° of AOM has been calculated using the *K*m(app) value for total ammonium (NH3 + NH4^+^) and not the *K*m(app) value for NH3 ^5,29^. Calculating the *a*° based on the *K*m(app) value for total ammonium allows for the *a*° of AOM to be compared with the *a*° of microorganisms that do not use NH3 as a sole energy generating substrate, such as ammonium assimilating heterotrophic bacteria or diatoms ^29^. While this is useful when evaluating competition for total ammonium in mixed communities or environmental settings, an *a*° calculated using the *K*m(app) value for NH3 may be more useful when directly comparing the interspecies competitiveness of AOM for the following reasons: i) the substrate for all AOM is NH3 and not NH4^+^ (see below) and ii) the *K*m(app) value for total ammonium is more dependent on the environmental factors it was measured at (e.g. pH, temperature, salinity) than the *K*m(app) for NH3.

All characterized AOA (with the exception of representatives of the genus *Nitrosocosmicus*) and the comammox bacterium *N. inopinata* possess much higher *a*° for total ammonium or NH3 (∼10 to 3000×) than the AOB, *N. oceani* or *N. europaea* (Fig. 2b,d), indicating that they are highly competitive in environments limited in either total ammonium or only NH3. However, due to the low number of published normalized *V*max values for AOB, *a*° could only be calculated for these two AOB representatives. Thus, extrapolations to the *a*° of all AOB species, based solely on these observations should be approached with caution.

The low variation in experimentally measured *V*max values (Fig. 2e) across all measured AOM in combination with the high variation in *K*m(app) values leads to a strong relationship between cellular *a*° and the reciprocal of *K*m(app) (Fig. 3) according to equation 2 (see Materials and Methods). AOM adapted to oligotrophic (low substrate) conditions should possess both a high substrate affinity (low *K*m(app)) and a high *a*^o 28^. Therefore, the AOM best suited for environments limited in total ammonium are the AOA belonging to the *Nitrosopumilales* and the comammox isolate *N. inopinata*, (top right corner of Fig. 3a). Overall, when looking at solely NH3 or total ammonium, the separation of species in these plots remains similar, with the exception that the acidophilic AOA belonging to the ‘*Ca.* Nitrosotaleales’ are predicted to be best suited for life in environments limited in NH3 (Fig. 3b). The adaptation correlates well with the fact the AOA ‘*Ca*. Nitrosotalea devanaterra’ Nd1 and ‘*Ca.* Nitrosotalea sinensis’ Nd2 were isolated from acidic soils with a pH of 4.5 and 4.7, respectively ^25, 45^, where NH3 is limiting even when total ammonium is not.

**Fig. 3.**
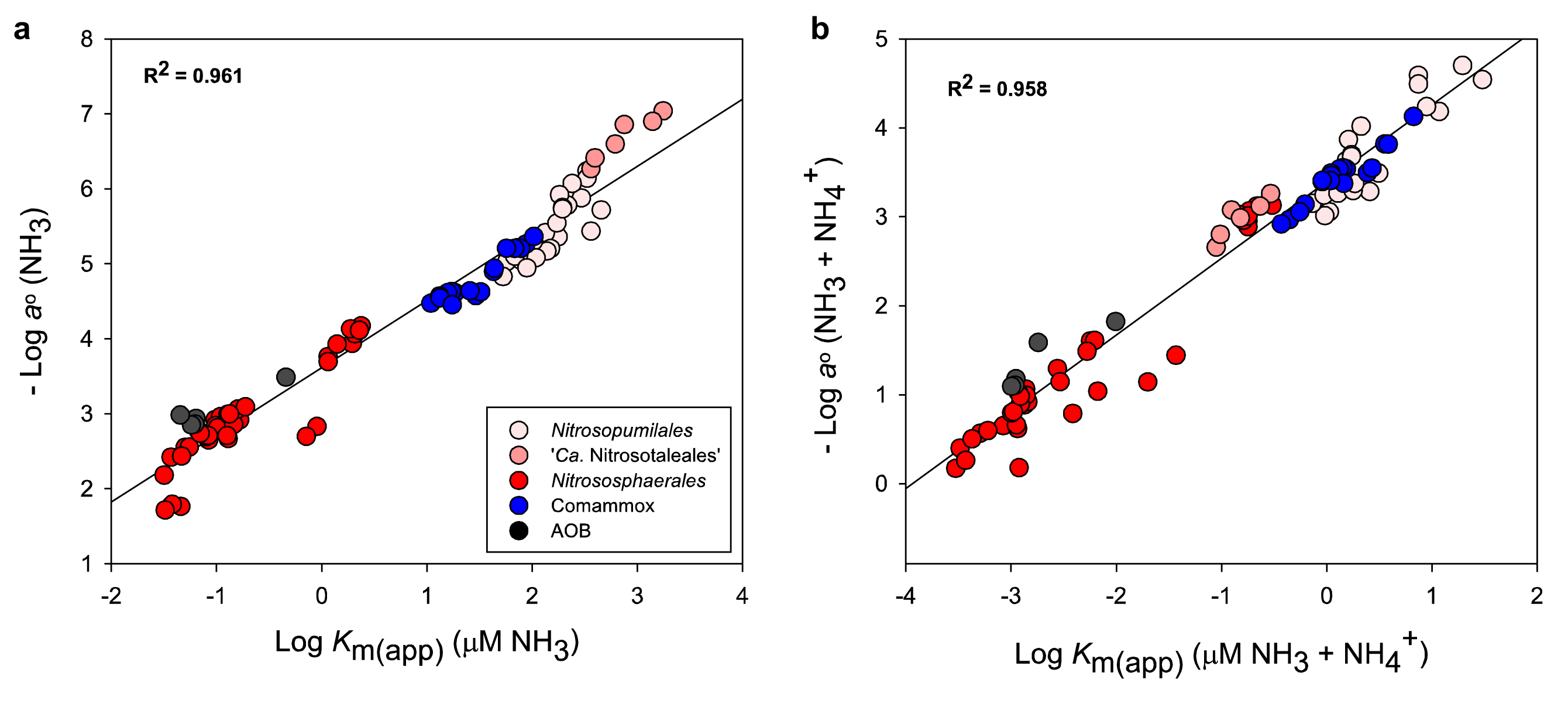
The reciprocal relationship between the substrate affinity (*K*_m(app))_ and specific substrate affinity (*a*°) of ammonia- oxidizing microorganisms (AOM). Reciprocal plots for both (a) total ammonium and (b) NH3 are depicted. The *K*m(app) and *a*° values correspond to the values presented for pure AOM isolates in Fig. 2. Data for AOA (red), comammox (blue), and AOB (black) are shown. The correlation (R^2^) indicates the linear relationship between the logarithmically transformed data points.

In either case, when looking at NH3 or total ammonium, the AOA belonging to the genus *Nitrosocosmicus* (‘*Ca.* N. oleophilus’ MY3 and ‘*Ca.* N. franklandus’ C13) and AOB populate the lower left section of these plots, indicating that they are not strong substrate competitors in NH3 or total ammonium limited environments (Fig. 3). If the cellular kinetic property of *V*max really is so similar across all AOB, AOA, and comammox species (Fig. 2e), then substrate competitiveness can be predicted from an AOMs *K*m(app) for either NH3 or total ammonium (Fig. 2a,c). This is especially helpful when characterizing enrichment cultures, where normalizing ammonia-oxidizing activity to cellular protein in order to obtain a comparable *V*max value is not possible. However, there is also a need for more kinetically characterized AOB and comammox species to confirm this hypothesis.

### The effect of environmental and cellular factors on AOA kinetic properties

The concentration of NH3 present in a particular growth medium or environment can vary by orders of magnitude, based solely on the pH, temperature, or salinity of the system ^54^. This is notable because at a given total ammonium concentration, the concentration of NH3 is ∼10 times higher at 70^°^C versus 30°C and ∼1000 times lower at pH 5.3 versus pH 8.4 (representative of maximum ranges tested). While it should be recognized that in our dataset no AOM were included that have a pH optimum between 5.3 and 7.0, the effect of pH and temperature on the ammonia oxidation kinetics of AOM must be considered in order to understand their ecophysiological niches. However, there was no correlation between the kinetic properties of AOM (*K*m(app), *V*max, and *a*°) measured in this study and their optimal growth temperature or pH. This lack of correlation between AOM species kinetic properties and growth conditions does not imply that the cellular kinetic properties of an individual AOM species will remain the same over a range of pH and temperature conditions. Therefore, we investigated the effect of pH and temperature variation on the substrate-dependent kinetic properties of the AOA strain ‘*Ca.* N. oleophilus’ MY3, and the effect of pH on the comammox strain *N. inopinata*.

### The effect of temperature

The effects of short-term temperature changes on the substrate-dependent kinetic properties of ‘*Ca.* N. oleophilus’ MY3 were determined. Temperature shifts of 5°C above and below the optimal growth temperature (30°C) had no effect on the *K*m(app) for total ammonium. However, the *K*m(app) for NH3, *V*max, and *a*° of ‘*Ca.* N. oleophilus’ MY3 all increased with increasing temperatures (Supplementary Fig. 6). Therefore, as temperature increased, ‘*Ca.* N. oleophilus’ MY3 displayed a lower substrate affinity (higher *K*m(app) for NH3) but would be able to turnover substrate with a higher *V*max and better compete for substrate with a higher *a*°. Increasing AOA *K*m(app) values for NH3 with increasing temperatures have also been observed across studies with *N. viennensis* EN76 (Supplementary Fig. 2). These cases are discussed in more detail in the Supplemental Results and Discussion. In addition, similar observations have previously been made for AOB strains belonging to the genus *Nitrosomonas* ^33, 34^. The increase in *V*max and *a*° can be explained in terms of the Van’t Hoff rule, which states that reaction velocity increases with temperature ^55^. While there is no theoretical upper limit to the reaction rate increase, the instability of proteins at elevated temperatures eventually becomes limiting.

The increase in *K*m(app) for NH3 (lower NH3 affinity) with increasing temperature is less straightforward to interpret. As this is a whole cell measurement, the observed differences may result from either broad cellular changes or from changes in individual enzymes involved in the ammonia oxidation pathway specifically. At the cellular level, changes in the proteinaceous surface layer (S-layer) or lipid cell membrane could affect substrate movement/transport and enzyme complex stability. It has been suggested that the negatively charged AOA S-layer proteins act as a substrate reservoir, trapping NH4^+^ and consequently increasing the NH3 concentration in the AOA pseudo-periplasmic space ^56^. It is interesting to note that sequenced representatives from the genus ‘*Ca*. Nitrosocosmicus’ lack the main S-layer protein (slp1) found in all *Nitrosopumilales*, *Nitrososphaerales*, and ‘*Ca*. Nitrosotaleales’ sequenced isolates ^52^. In addition, it has been demonstrated that elevated temperatures significantly alter the lipid composition in the AOA cell membrane ^57, 58^. However, it is unclear how changes in the S-layer or cell membrane affect whole cell substrate affinity. On the single enzyme level, previous studies have shown the same trend of decreasing substrate affinity and increasing maximal reaction velocity with increasing temperatures, due to altered protein structures and an increased enzyme-substrate dissociation constant ^59, 60^.

Notably, differing optimum growth and activity conditions were previously determined for the marine AOB strain *Nitrosomonas cryotolerans* ^34^. These observations raise interesting, albeit unanswered, questions about why the growth and activity temperature optima are or can be uncoupled in AOM, and what this means for AOM niche differentiation and their competitiveness *in-situ*. Moving forward, investigations into the growth and cellular kinetic properties of AOM across a range of environmental factor gradients will be essential in understanding competition between AOM in engineered and environmental systems.

### The effect of pH

The effects of short-term pH changes on the substrate-dependent kinetics of ‘*Ca.* N. oleophilus’ MY3 and *N. inopinata* were determined. The *V*max of both ‘*Ca*. N. oleophilus MY3’ and *N. inopinata* were stable at 37.3 ± 6.6 μmol N mg protein^-1^ h^-1^ and 11.2 ± 2.5 μmol N mg protein^-1^ h^-1^, respectively, in medium with a pH between ∼6.5 to ∼8.5 (Supplementary Table 3). However, the *K*m(app) for total ammonium of ‘*Ca*. N. oleophilus MY3’ and *N. inopinata* decreased by more than an order of magnitude (∼11×) across this pH range, while the *K*m(app) for NH3 remained more stable, increasing only 3 to 4 times (Fig. 4). As an increase of pH shifts the NH3 + NH4^+^ equilibrium towards NH3, this suggests that the actual substrate used by AOA and comammox is indeed the undissociated form (NH3) rather than the ammonium ion (NH_4_ ^+^), as previously demonstrated for AOB ^34, 35, 51, 61^. Interestingly, the only exception to this rule to date is the gammaproteobacterial marine AOB *Nitrosococcus oceani*. The reported *K*m(app) for total ammonium of *N. oceani* remained more stable (∼2.3×) than the *K*m(app) for NH3 (78×) when the pH was shifted from 6.3-8.6 ^62^. With this exception in mind, it is likely that all AOA, AOB, and comammox compete for NH3 as their substrate in the environment.

**Fig. 4.**
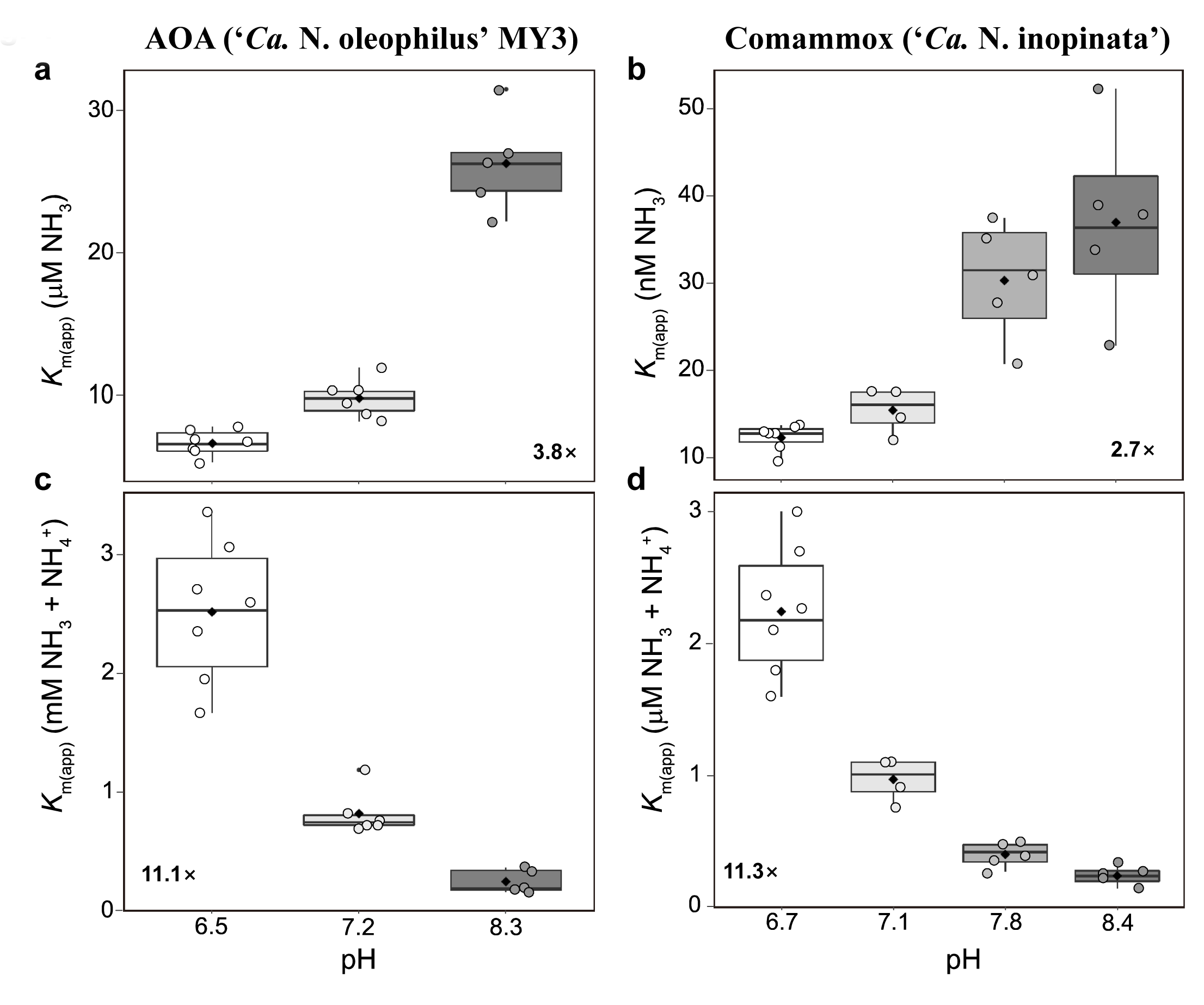
The effect of medium pH on the substrate affinity of ‘*Ca*. N. oleophilus MY3’ and *N. inopinata*. The substrate affinities for both (a,b) NH3 and (c,d) total ammonium (NH3 + NH4^+^) are provided. Individual substrate affinity values determined at each pH are shown as single points (circles). The boxes represent the first and third quartiles (25% to 75%) of the substrate affinity range under each condition. The median (line within the boxes) and mean substrate affinity (black diamonds) values are also indicated. The whiskers represent the most extreme values within 1.58× of quartile range. The variation of the substrate affinity values across the entire tested pH range are indicated in each panel. In all four instances there was a significant difference between the affinity at the lowest pH and the highest pH, as determined by a Student’s t-test (*p*<0.005). The average substrate affinity values for ‘*Ca.* N. oleophilus MY3’ and *N. inopinata* at each pH are provided in Supplementary Table 3.

It is important to note that the substrate affinities reported here represent whole cell affinities and not the substrate affinity of ammonia monooxygenase (AMO) enzymes. Therefore, further experimental investigation with purified AMO and ammonia/ammonium transporter proteins is warranted. Although NH3 can freely diffuse passively into AOM, this does not mean that the cellular affinity reported here is necessarily unrelated to the transporter-mediated movement of NH3/NH4^+^ into AOM cells. For example, AOB have previously been shown to accumulate very high (1 M) intracellular NH4^+^ concentrations ^63^. This high intracellular NH4^+^ concentration may provide a concentrated substrate reservoir, indirectly increasing the concentration of NH3 around the AMO enzyme complex. However, it is unknown if such a concentration mechanism would be more important for an AOB with a low substrate affinity (e.g. *N. europaea*) or for an AOA living in extremely substrate-limited environments (e.g. *N. maritimus*).

### The effect of cell morphology

All AOM share the primary enzyme involved in ammonia oxidation, AMO, which is located in the cytoplasmic membrane with its substrate-binding site most likely facing the outside of the cell ^56^. Therefore, a higher cellular surface area to volume (SA/V) ratio likely contributes to an increase in *a*°, as it increases the space available for AMO and the chance to bind NH3 at very low concentrations. This assumption is based on the hypothesis that an increased abundance of uptake enzymes (e.g., permeases) leads to a higher *a*^o 28, 64^. In fact, the SA/V ratio of AOM (Supplementary Table 4) correlates to the log of their observed *K*m(app) for NH3 (R^2^ = 0.82), *K*m(app) for total ammonium (R^2^ = 0.74), *a*° for NH3 (R^2^ = 0.77), and *a*° for total ammonium (R^2^ = 0.81; Fig. 5). Therefore, the SA/V ratio of newly cultured AOM might be a useful indicator for these cellular kinetic properties. Consequently, AOM with a high SA/V ratio will likely outcompete other AOM in many natural aquatic and terrestrial environments, such as the pelagic marine water column that has a very low standing total ammonium pool. Consistently, these oligotrophic environments have already been postulated to select for organisms with a high SA/V ratio, enhancing their nutrient uptake capabilities ^65, 66^.

**Fig. 5.**
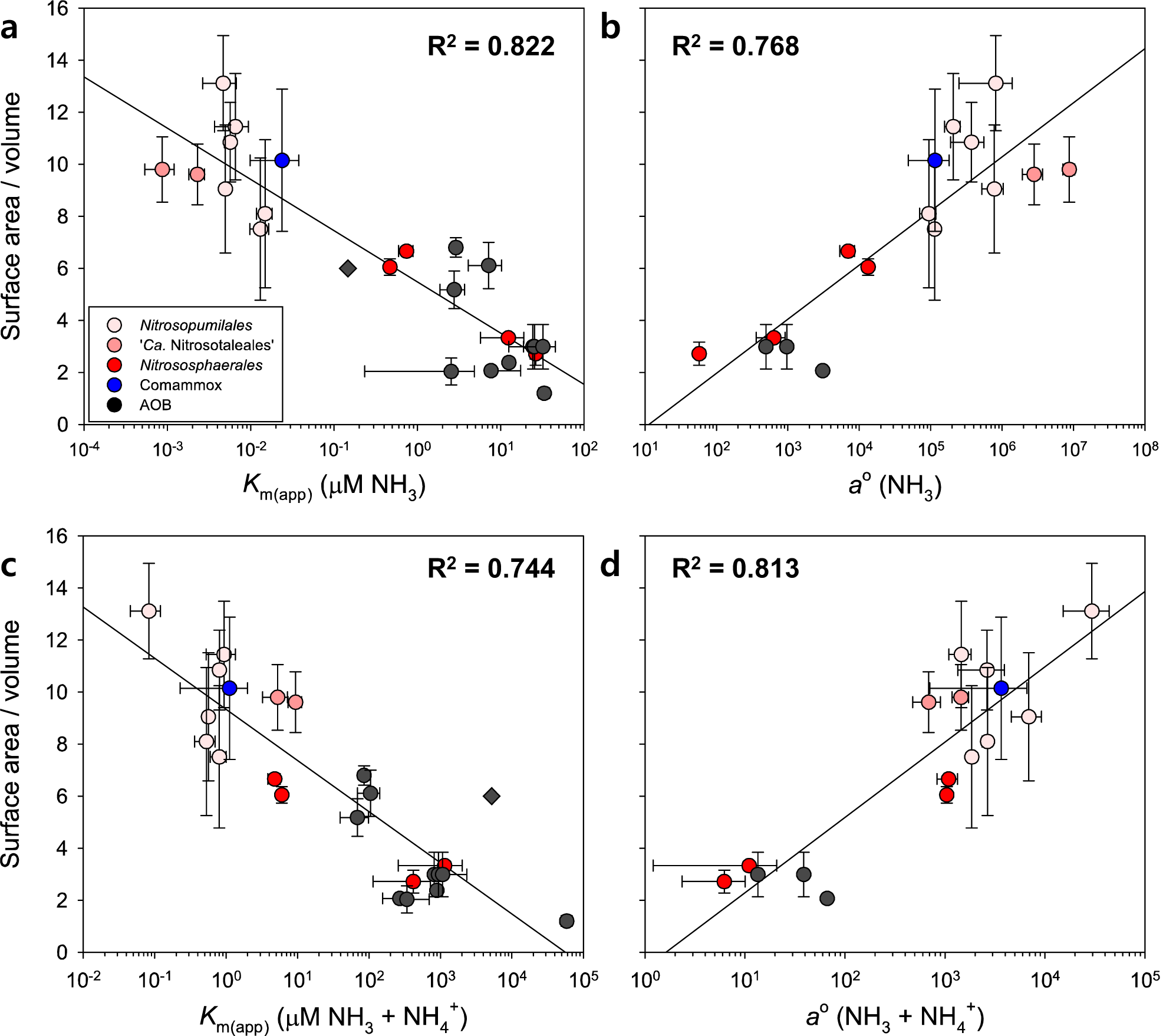
Logarithmic correlation of (a) substrate affinity (*K*_m(app))_ and (b) specific substrate affinity (*a*°) with the cellular surface area to volume ratio of ammonia-oxidizing microorganisms (AOM). All *K*m(app) and *a*° values correspond to values presented in Fig. 2. The surface area to volume (SA/V) ratio calculations for each AOM are provided in Supplementary Table 4. Data for AOA (red), comammox (blue), and AOB (black) are shown. The three different gradations of red differentiate three distinct AOA phylogenetic lineages. The error bars represent the standard deviation of replicate kinetic experiments or SA/V ratio measurements of each AOM strain. The logarithmic correlation (R^2^) value was calculated from the average values of each AOM and is presented on a semi-log axis.

The correlation between the SA/V ratio and cellular kinetic properties of AOM sheds some light on the unusual kinetic properties of the AOA belonging to the genus *Nitrosocosmicus*. Both ‘*Ca.* N. oleophilus’ MY3 and ‘*Ca.* N. franklandus’ C13 possess a very low SA/V ratio compared to other AOA isolates and they both possess several characteristics normally associated with AOB – high substrate tolerances ^48–50^, low affinities for NH3, and a low *a*° for NH3 – that are not consistent with the long-held convention that all AOA are much stronger competitors for NH3 than AOB in substrate-limited environments. Therefore, the individual cell morphology of AOM may have a direct relationship with their cellular kinetic properties. Although this is only a correlation-based observation, it highlights that further investigation into these characteristics is warranted.

In addition to cellular morphology, the size of cell aggregates can affect the kinetic properties of AOM ^67^. Cell aggregates have a lower SA/V ratio than individual cells, which can decrease diffusion rates and create microscale substrate/oxygen gradients within aggregates ^68^. In order to ensure that the large differences in substrate affinity among AOA are not caused by differences in cell aggregation, the aggregate size of ‘*Ca.* N. uzonensis’ N4, ‘*Ca.* N. oleophilus’ MY3, and *N. piranensis* D3C cultures were inspected before and after MR experiments (Supplementary Fig. 7). No aggregation pattern was observed that would explain the multiple orders of magnitude differences in substrate affinity between these AOA. In fact, of the three AOA investigated, the only strain to form large cell aggregates either before or after MR experiments was *N. piranensis* D3C, which has one of the highest measured substrate affinities (lowest *K*m(app) for NH3). In contrast, the cell aggregate size of ‘*Ca.* N. oleophilus’ MY3 and ‘*Ca.* N. uzonensis’ N4 were unaffected by the MR experiment and remained relatively small (Supplementary Fig. 7). As ‘*Ca.* N. oleophilus’ MY3 has one of the lowest substrate affinities (highest *K*m(app) for NH3) and formed only small cell aggregates, the low substrate affinity of ‘*Ca.* N. oleophilus’ MY3 was not an artefact caused by cell aggregation.

Taken together, both environmental (pH and temperature) and AOM cellular (SA/V ratio) factors affect or are related to the observable cellular kinetic properties of individual AOM species. These factors need to be considered when investigating AOM competition or niche differentiation *in-situ*, as they are often in flux in environmental settings. This can be especially true considering cell morphology, which is often dependent on growth conditions ^69^. However, the plasticity of the cellular kinetic properties within individual AOM species does not explain the larger trends observed here across AOA lineages or between AOM.

### Concluding remarks

In this study we substantially extended the set of available substrate oxidation kinetic parameters for AOA by the analysis of pure cultures or enrichments from various lineages within this guild.

Furthermore, our kinetic data obtained at different pH values supports the hypothesis that, like for AOB, the substrate for AOA and comammox is NH3. Together, our findings provide novel insights for our understanding of niche differentiation among AOM and demonstrate a surprising variability of the inferred kinetic parameters among AOA. Thus, the long-standing and recently questioned^5^ hypothesis that all AOA have extremely high substrate affinities and specific substrate affinities, can no longer be maintained. The observed links between AOA kinetic properties, phylogeny, and cell morphology also enables the formulation of testable hypotheses on nitrification kinetics in systems thus far characterized solely with molecular (e.g. amplicon sequencing or metagenomic) tools.

As environmental factors such as temperature and pH influence kinetic parameters of AOA including their cellular affinity for NH3, future analyses of kinetic parameters of AOM should not only be performed at their optimal growth conditions, but also over a range of conditions that reflect their environmental niches. Such experiments will generate a more informative picture on AOM competition and niche differentiation.

### Newly isolated Nitrosotenuis species

The isolated strain N4 is a novel species of the genus *Nitrosotenuis* of the order *Nitrosopumilales*, and we propose the following candidate status:

#### Taxonomy

**(i) Etymology.** The taxonomy for ‘*Candidatus* Nitrosotenuis uzonensis’ sp. nov. is as follows: Nitrosus (Latin masculine adjective), nitrous; tenuis (Latin masculine adjective), small/slender; uzonensis (Latin neutrum genitive), from Uzon.
**(ii) Locality.** A terrestrial thermal spring located in the Uzon caldera on the Kamchatka peninsula, Russia.
**(iii) Diagnosis.** A chemolithoautotrophic ammonia oxidizer of the phylum Thaumarchaeota, which is straight and rod-shaped, with a diameter of 0.2-0.3 µ m and a length of 0.4-1.7 µ m. Growth over a period of several years has been maintained in a medium with a pH of 7.5 at 37°C. It belongs to the AOA order *Nitrosopumilales* (group I.1a). AOA with almost identical 16S rRNA and *amoA* gene sequences have been detected in various environments, including soil and groundwater^22, 40, 41^.

## Materials and Methods

### Cultivation of ammonia oxidizers

Several previously described growth media were used to cultivate the AOM used in this study. A comprehensive guide with medium components and cultivation conditions is provided in the Supplementary Materials and Methods, Supplementary Tables 1, and 2. Briefly, all cultures were grown without shaking, in the dark, at their optimum growth temperature and pH, unless otherwise stated. Ammonium (NH4Cl) from pre-sterilized stocks was added as substrate as needed. The growth medium of *Nitrosarchaeum koreense* MY1, ‘*Ca.* Nitrosotenuis chungbukensis’ MY2, ‘*Ca.* Nitrosotenuis uzonensis’ N4, *N. maritimus* SCM1, *Nitrosopumilus piranensis* D3C, and *Nitrosopumilus adriaticus* NF5 was supplemented with sodium pyruvate (0.5 mM) at all times. The pH of all growth media were adjusted when necessary by addition of sterile NaHCO3. Ammonia oxidation activity was determined by measuring ammonium, nitrite, and nitrate concentrations photometrically ^70–73^ using an Infinite 200 Pro M Nano+ spectrophotometer (Tecan Group AG, Switzerland).

### Novel AOA enrichments and pure culture

The sampling site, enrichment process, and initial strain characterization details for the two novel thermophilic AOA enrichment cultures ‘*Ca*. Nitrosofervidus tenchongensis’ DRC1, and ‘*Ca*. Nitrososphaera nevadensis’ GerE used in this study are provided in the Supplementary Materials and Methods. In addition, in this study, ‘*Ca*. N. uzonensis’ N4 was isolated as a pure culture from a previously described geothermal spring enrichment culture ^41^. Further details are provided below and in the Supplementary Materials and Methods.

### Phylogenetic analysis

A phylogenetic tree for all strains used in this study as well as select reference strains were calculated using IQ-TREE v 1.6.2 ^74^. Automatic model determination using modelFinder ^75^ identified LG+F+R5 as the best-fit model according to the Bayesian Information Criterion (BIC). Bipartition support was determined with ultrafast bootstraps (UFboots ^76^). The tree was based on an alignment of 34 universal genes (43 markers) extracted from genomes, aligned and concatenated using CheckM ^77^.

### Substrate-dependent oxygen uptake measurements

Cellular substrate oxidation kinetics were determined from instantaneous substrate-dependent oxygen uptake measurements as previously described ^5,29, 78^. Briefly, measurements were performed with a microrespirometry (MR) system, equipped with a PA 2000 picoammeter and a 500 μm tip diameter OX-MR oxygen microsensor (Unisense, Denmark), polarized continuously for at least 24 hours before use.

Active AOA, AOB, and *N. inopinata* cells were harvested (4000×*g*, 10 min, 20^°^C) from ammonium replete active cultures, using 10 kDa-cutoff, Amicon Ultra-15 centrifugal filter units (Merck Millipore, Germany). Concentrated cells were washed with and resuspended in substrate-free medium appropriate for the respective cultures. Exceptions were ‘*Ca.* Nitrosocosmicus franklandus’ C13 and the marine AOA, *N. maritimus* SCM1, *N. piranensis* D3C, and *N. adriaticus* NF5. These four AOA strains were not active in the MR chambers after attempts to concentrate their biomass. Therefore, ammonium concentrations were monitored daily for these four cultures, and cells were used without concentration for MR promptly upon substrate depletion ^29^. AOM harvested cells for MR experiments were incubated for at least 30 min in a recirculating water bath set to the experimental temperature (Supplementary Tables 2 and 3) prior to being transferred to the MR chambers (∼2 ml).

In addition to MR experiments at optimal growth temperature and pH (Supplementary Table 2), MR experiments were also performed at non optimal growth temperatures and medium pH (Supplementary Table 3). ‘*Ca.* N. oleophilus’ MY3 was cultivated at 30^°^C, harvested with centrifugal filter units (see above), and incubated for ∼2 hours in substrate-free medium across a range of temperatures (25, 30, and 35^°^C). MR experiments were then performed at the respective preincubation temperature. Likewise, *N. inopinata* and ‘*Ca.* N. oleophilus’ MY3 cells were harvested with centrifugal filter units (see above) and resuspended in substrate-free medium containing 10 mM HEPES (pH 7.4). The pH was adjusted to 6.5-8.4 with 1 M HCl or 1 M NaOH (Supplementary Table 3). These cultures were then incubated at their optimum growth temperature for ∼1 hour prior to cellular kinetic measurements. Culture pH was determined before and after oxygen uptake measurements to confirm the pH did not change during MR. Substrate-dependent oxygen uptake measurements were performed as described below.

For all MR experiments, glass MR chambers containing glass-coated magnetic stir bars were filled headspace-free, sealed with MR injection lids, and submerged in a recirculating water bath. An OX-MR microsensor was inserted into each MR chamber and left to equilibrate (300 rpm, ∼1 h). Exact temperatures used for each culture and experiment are provided in Tables S2 and S3. Stable background sensor signal drift was measured for at least 15 min prior to initial substrate injections, and the background oxygen consumption rate was subtracted from the measured oxygen uptake rates. Hamilton syringes (10 or 50 μl; Hamilton, USA) were used to inject NH4Cl stock solutions into MR chambers. Both single and multiple trace oxygen uptake measurements were performed. For single trace measurements, a single substrate injection was performed, and oxygen uptake was recorded until substrate depletion. For multiple trace measurements, multiple injections of varying substrate concentration were performed in a single MR chamber. Once stable, discrete slopes of oxygen uptake were calculated following each substrate injection. Immediately following oxygen uptake measurements, the total ammonium concentration and pH of the MR chamber contents were determined. The cells were stored at -20^°^C for protein analysis. Cells were lysed with the Bacterial Protein Extraction Reagent (BPER, Thermo Scientific) and the total protein content was determined photometrically with the Pierce bicinchoninic acid (BCA) Protein Assay Kit (Thermo Scientific) as per the manufacturer’s instructions. Before and after MR assays of *N. piranensis* D3C, ‘*Ca.* N. uzonensis’ N4, and ‘*Ca.* N. oleophilus’ MY3, an aliquot of culture was filtered onto membranes (0.2 µ m polycarbonate GTTP membranes; Merck Milipore, Germany) and DAPI (4’,6-diamidino-2-phenylindole; 10 µ g ml^-1^; 5 min; room temperature) stained prior to microscopic measurement of biomass cell aggregate size, as described previously ^67, 79^.

### Calculation of kinetic properties

*K*m(app) and *V*max were calculated from both single and multiple trace substrate-dependent oxygen uptake measurements. Total ammonium (NH3 + NH4^+^) oxidation rates were calculated from oxygen uptake measurements using a substrate to oxygen consumption ratio of 1:1.5 ^5,29^. Total ammonium uptake rates were fitted to a Michaelis-Menten model using the equation:

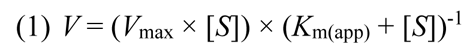

where *V* is the reaction rate (μM h^-1^), *V*max is the maximum reaction rate (μM h^-1^), *S* is the total ammonium concentration (μM), and *K*m(app) is the reaction half saturation concentration (μM). A nonlinear least squares regression analysis was used to estimate *K*m(app) and *V*max ^80^. The *K*m(app) for NH3 for each strain was calculated based on the *K*m(app) for total ammonium, incubation temperature, pH, and salinity ^81^. *K*m(app) values for AOM not determined in this study were compiled from the literature ^5,29, 33, 34, 42, 51, 82–84^. If only total ammonium information was given by the authors for *K*m(app), the corresponding NH3 values were calculated based on the reported experimental temperature, pH, and salinity values. *V*max values of pure cultures were normalized to culture protein content. The specific substrate affinity (*a*°; l g wet cells^-1^ h^-1^) of each pure culture strain was calculated using the equation:

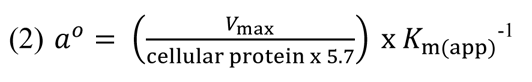

Where the *V*max is normalized to the protein concentration (g l^-1^) of the culture in the MR chamber and the factor of 5.7 g wet cell weight per g of protein was used for all AOM ^5, 29, 64^. The *a*° for NH3 or total ammonium were calculated using the respective *K*m(app) for NH3 or total ammonium.

#### Cell surface area to volume ratio calculation

Approximate cell surface area to volume ratios were determined using cell dimensions provided by or calculated from previously published phase contrast, transmission electron, or scanning electron microscopy images (Supplementary Table 4). The following equations for the surface (SA) area and volume (V) of a sphere (3) and rod (4) were used:

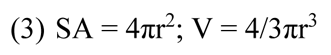

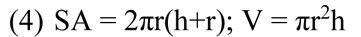

where r is the cell radius (µ m) and h is the cell length (µ m). The cell size and volume from published phase contrast images were verified using MicrobeTracker ^85^.

## Acknowledgements

We would like to thank Márton Palatinszky, Ping Han, Michael Lukumbuzya, and Dimitra Sakoula for their assistance with microscopy and culture maintenance. CJS and DK were supported by the Wittgenstein award of the Austrian Science Fund FWF (Z383-B) to MW. PP and CJS were supported by the Austrian Science Fund FWF through the Young Investigators Research Grant program (ZK76). M-Y Jung was supported by the Research Institute for Basic Sciences (RIBS) of Jeju National University through the National Research Foundation of Korea (NRF) grant funded by the Ministry of Education (2019R1A6A1A10072987) and NRF grant funded by the Korea government (MSIT) (NRF-2021R1C1C1008303). CW was supported by a University of East Anglia-funded PhD studentship. LLM was supported by a Royal Society Dorothy Hodgkin Research Fellowship (DH150187) and by a European Research Council (ERC) Starting Grant (UNITY 852993).

## Author contributions

M-YJ, CJS HD, and MW designed the study and wrote the manuscript with the help of all authors. M-YJ and CJS performed the kinetic experiments with the help of KDK, LH, BB, L L-M, and CW. Additional data analysis was performed by AM, S-KR, PP, GWN, JRT, and CWH.

