## Supplemental text, tables, and figures for "Ammonia-oxidizing archaea possess a wide range of cellular ammonia affinities"

**This document includes:**

Supplementary Material and Methods

Supplementary Results and Discussion

Supplementary Tables S1 to S4

Supplementary Figures S1 to S8

Supplementary References

### Supplementary Materials and Methods

#### *Isolation of* **'Candidatus *Nitrosotenuis uzonensis*' *N4***

'*Ca.* N. uzonensis' N4 was isolated from a previously described thermal spring ammonia-oxidizing enrichment culture [1]. Further enrichment and subsequent purification of '*Ca*. N. uzonensis N4' was performed over the course of 6 years. Successive cultures were maintained in a mineral salt medium (Supplementary Table 1) containing a mixture of antibiotics (ampicillin and penicillin-G; 50 μg $ml^{-1}$). Cultures were transferred approximately every three weeks, and a pure culture designated strain N4 was obtained through filtration (0.22-μm syringe filter; Sigma Aldrich) and subsequent end-point serial dilution in deep well 96-well plates containing medium supplemented with ammonium chloride (100 μM) and pyruvate (50 μM). The activity of strain N4 was monitored by determining ammonium consumption and nitrite production photometrically [2-5]. After isolation, '*Ca*. N. uzonensis' N4 was routinely grown in pyruvate supplemented medium without antibiotics. Cultures were maintained in the dark, without shaking at 37℃. A general bacterial 16S rRNA gene PCR (27F and 1492R) [6] was used to verify the absence of any bacterial contaminants as described in Weisburg et al.,[7]. Culture purity was confirmed and routinely monitored by fluorescence *in situ* hybridization (FISH) as previously described [8] with a Cy3-labeled *Archaea*-specific probe (Arc915) [9] and a FAM-labeled *Bacteria*-specific probe (EUB338) [10]. To visualize all cells, 4',6-diamidino-2-phenylindole (DAPI) staining was used. In addition, the absence of bacterial growth in three nutrient-rich media broths was routinely observed (i.e., lysogeny broth, Reasoners 2A broth, and tryptic soy broth) [11].

#### *New thermophilic AOA enrichment cultures*

Cultivation of '*Ca.* Nitrosocaldus yellowstonensis' HL72 (formerly published as '*Ca.* N. yelllowstonii' HL72) from hot spring sediment from Yellowstone National Park, Wyoming (USA) was previously described [12]. Two additional thermophilic AOA cultures were enriched using the same medium and approach used to cultivate HL72. '*Ca*. Nitrosofervidus tenchongensis' DRC1 was cultivated at 72˚C from the Direchi Geothermal Pond (地热池) in the Rehai geothermal field in Tengchong, Yunnan Province, China. The temperature and pH at the sampling site (N 24.95009°, E 98.43807°) were 83˚C and pH 8.3, respectively. Additional geochemical characterization of the site has been described elsewhere [13,14]. '*Ca*. Nitrososphaera nevadensis' GerE is a moderately thermophilic strain growing at 50˚C and was cultivated from a low temperature spring (45˚C, pH 8.0) located adjacent to Great Boiling Spring (N 40.66148˚, E 119.36613˚) in the Great Basin of Nevada (USA) [15].

***Cultivation conditions of ammonia-oxidizing microorganisms***

The medium description and cultivation conditions of all ammonia-oxidizing microorganisms (AOM) used in this study are provided in Supplementary Tables 1 and 2. Briefly, all AOM cultures were grown in Schott bottles at their optimal temperature and pH, in the dark, without shaking, unless otherwise noted. When necessary, the initial pH of all AOM media was adjusted with 1 M HCl and 1 M NaOH. Substrate was provided in the form of $NH_4Cl$ from pre-sterilized stocks and replenished as necessary. Sterile sodium bicarbonate (1M) was added as necessary in order to adjust culture pH. An initial concentration of 0.5 mM sodium pyruvate was added to the cultivation medium of *N. koreense* MY1, '*Ca.* N. chungbukensis' MY2, '*Ca.* N. uzonensis' N4, *N. maritimus* SCM1, *N. piranensis* D3C, and *N. adriaticus* NF5 as a hydrogen peroxide ($H_2O_2$) scavenger [16].

***Nitrite-dependent ammonia oxidation kinetics***

In addition to microrespirometry measurements, ammonia oxidation kinetics of '*Ca*. Nitrosocosmicus franklandus' C13 were also determined by nitrite accumulation activity assays performed with concentrated cell suspensions as previously described [17]. Briefly, '*Ca*. Nitrosocosmicus franklandus' C13 was grown in batch culture until mid-exponential growth phase was reached (corresponding to 600-700 μM nitrite accumulated in the medium). Cells from 1.6 L of culture were harvested onto 0.22 μm pore size, 45 mm diameter polyethersulfone (PES) filter (PALL). Cells were washed with 10 mM HEPES (pH 7.3)-buffered medium with no added ammonium to remove residual ammonium and nitrite. Cells were then resuspended in 200 mL 10 mM HEPES (pH 7.5)-buffered medium with no added ammonium. Concentrated cells were incubated at 37°C without ammonium for 60 min to allow the endogenous respiration to cease. For cell counts, cells were stained with DAPI and counted on 0.22 μm pore size black polycarbonate filters using fluorescence microscopy as previously described [18].

For the kinetics assays, cell concentrations were adjusted to approximately $1 \times 10^7$ $ml^{-1}$ to ensure that the rates of nitrite accumulation were always linear over the course of the entire assay. Aliquots of 5 ml cell suspension were added to acid-washed 23 ml glass vials which were sealed with twice autoclaved grey butyl rubber septa. The assays were performed in a 37°C or 42°C static water bath and all treatments were performed in triplicate vials. Ammonia-oxidizing activity was initiated by adding $NH_4Cl$ at the following final concentrations: 5, 10, 25, 50, 100, 250, 500, and 1000 μM. 100 μl volume was removed from each vial at 15 min intervals using a needle and syringe, and nitrite concentration was measured immediately after sampling. Assays were run over a period of 60 min. $K_{m(app)}$ and $V_{max}$ values were determined using the Hyper32 kinetics package

(Hyper32.exe, version 1.0.0, 2003). Protein determination was carried out using the PierceTM BCA Protein Assay Kit (ThermoFisher Scientific) according to the manufacturer's instructions.

Nitrite concentrations were determined colorimetrically using the Griess reagent in 96-well flat bottom clear microtiter plates as previously described [18]. Briefly, 20 µl sulphanilamide solution (5 g $L^{-1}$ in 2.4 M HCl) was added to 100 µl of sample or standard, followed by the addition 20 µl of naphthylethylenediamide solution (3 g $L^{-1}$ in 0.12 M HCl). Standards were performed in duplicate, prepared using $KNO_2$ and ranging from 0.5 to 50 µM. Absorbance was recorded at a wavelength of 540 nm using a VersaMax platereader (Molecular Devices, CA, US).

***Statistical analysis***

Statistical analyses were performed using the R environment for statistical computing (http://www.R-project.org/) and SigmaPlot 11.0 (Systat Software Inc., San Jose, CA, USA). An independent-sample t-test was performed to assess the significant difference between two groups, and *p* values less than 0.05 were considered to be statistically significant.

### Supplementary Results and Discussion

***Cellular kinetic properties across studies***

**The effect of differential medium conditions on AOM cellular kinetic properties.**

Differences in growth medium composition may affect the physiological state of AOM, potentially leading to unreproducible cellular kinetic measurements. To test this hypothesis, the cellular kinetic properties of *N. inopinata* were measured under optimal growth temperature and pH, in a different growth medium than used previously for the kinetic characterization of this organism [19]. In these experiments we determined only a minute effect on the $K_{m(app)}$ for $NH_3$, $K_{m(app)}$ for total ammonium ($NH_3$ + $NH_4^+$), or $V_{max}$, and no significant effect on the $a^o$ for $NH_3$ or total ammonium ($NH_3$ + $NH_4^+$) ($p > 0.7$) (Fig. 2, Supplementary Figs. 2 and 8). In addition, the substrate affinity for $NH_3$ of *Nitrosomonas europaea* ($K_{m(app)}$=23.3 µM, s.d.=1.5; Supplementary Fig. 8) and *Nitrosarchaeum koreense* MY1 ($K_{m(app)}$=4.8 nM, s.d.=0.5; Supplementary Figs. 1 and 2) which were previously determined in different media, were confirmed in this study. Notably, there was no significant difference ($p$ = 0.74) between the cellular kinetics of *N. koreense* MY1 when determined as an enrichment culture (containing <10% partner bacteria) [20] or as pure culture in this study (Supplementary Fig. 2). This indicates that the different growth media used to culture AOM in this study, does not affect the reproducibly of the cellular kinetic property measurements.

**Activity versus growth kinetic properties**

The substrate affinity of *N. maritimus* SCM1 for $NH_3$ was reported to be ~3 nM, as determined by substrate-dependent oxygen uptake measurements, which was about three orders of magnitude higher than the affinity for $NH_3$ of any characterized ammonia-oxidizing bacterium (AOB) [21]. Based on this analysis of *N. maritimus*, it was widely generalized in many studies that AOA possess a high affinity for $NH_3$, and interpreted as an explanation for the high abundance of AOA compared with AOB in nutrient poor environments [22] until it was determined that non-marine AOA do not all possess such a high affinity for $NH_3$ [19].

Recently, the primary report of the substrate affinity of *N. maritimus* SCM1 for $NH_3$ [21] was called into question [23], and the revised substrate affinity of *N. maritimus* SCM1 was reported to be ~2 µM. This represents a substrate affinity three orders of magnitude lower than previously determined, and within the same range as the substrate affinity of the AOB *Nitrosomonas europaea* - an AOB adapted to high substrate concentrations [23]. Both sets of kinetic experiments were performed in the same growth medium and at the same temperature (30℃). In addition, the strains used displayed near-identical maximum specific growth rates (0.027 versus 0.028 $h^{-1}$) in batch culture [23]. Finding an explanation of these sharply contrasting observations is important, not just for defining the kinetic characteristics of *N. maritimus* SCM1, but for how widely applicable and reproducible these types of kinetic studies are.

It was speculated that *N. maritimus* SCM1 may have become domesticated and lost its high substrate affinity during years of laboratory cultivation at an increased substrate concentration, as the selective pressure which it was under in the environment was removed. To test this hypothesis, we investigated the kinetic properties of the *N. maritimus* SCM1 strain used in Hink *et al.* [23] (kindly provided by the lab of James I. Prosser), using the microrespirometry (MR) method that was applied in Martens-Habbena *et al.* [21]. As we were able to reproduce the substrate affinity values for *N. maritimus* SCM1 ($K_{m(app)}$ = ~3.33 nM for $NH_3$) reported by Martens-Habbena *et al.* [21], we can rule out domestication of *N. maritimus* SCM1 as the cause of these contrasting results (Fig. 2 and Supplementary Figs. 1 and 2).

Therefore, we propose that a possible explanation of the observed contrasting results represents the difference between the half saturation activity/growth properties ($K_m$/$K_s$) as determined by Michaelis-Menten (S1) and Monod kinetics (S2), respectively:

$$\text{(S1)} \quad V = (V_{max} \times [S]) \times (K_{m(app)} + [S])^{-1}$$

$$\text{(S2)} \quad \mu = (\mu_{max} \times [S]) \times (K_{s(app)} + [S])^{-1}$$

Although both equations utilize the same form, the Michaelis-Menten equation model activity whereas the Monod equation models growth. This represents the difference between the $K_m$ (the half saturation of activity, or the substrate concentration when the activity rate is half of the maximal activity rate) and the $K_s$ (the half saturation of growth, or the substrate concentration when the growth rate is half of the maximal growth rate). The major difference is that cellular processes involved in microbial growth such as repair, turnover, stress response, and division are accounted for in the Monod but not in the Michaelis-Menten model [24]. With this rationale, the $K_{m(app)}$ reported in the growth experiment in Hink *et al.* [23] may actually represent a $K_{s(app)}$ value, compared with the $K_{m(app)}$ activity values generated in this study and by Martens-Habbena *et al.* [21]. This explanation has recently also been proposed to describe these observed differences, however without experimentally ruling out strain domestication [25]. Even with this explanation, it is still hard to explain why these two kinetic parameters ($K_{m(app)}$ and $K_{s(app)}$) observed under ideal conditions vary by three orders of magnitude.

**The effect of culture growth stage and temperature on the cellular kinetics of AOM**

The cellular kinetics of the enrichment culture '*Ca.* N. uzonensis' N4 have previously been reported[19]. Since this report, '*Ca.* N. uzonensis' N4 has been isolated (detailed above) and the cellular kinetics were re-determined here using the pure '*Ca.* N. uzonensis' N4 culture.

Surprisingly, the substrate affinity of the pure '*Ca.* N. uzonensis' N4 culture was orders of magnitude higher (lower $K_{m(app)}$) than what was originally measured for the enrichment culture (Supplementary Figs. 1 and 2). Notably, the stoichiometry of $NH_4^+$ and $O_2$ consumption was close to 1:1.5 (1:1.48, s.d=0.02, *n*=5) in the enrichment culture when measured in 2017. Therefore, a strong microrespirometry bias caused by respiration of heterotrophic contaminants had been ruled out.

One possible explanation for the different results between enrichment and pure culture characterization is the difference in the growth phase of the cultures before the MR experiments. In Kits *et al.* [19] '*Ca.* N. uzonensis' N4 enrichment cultures were harvested from batch cultures at the onset of substrate depletion (onset of stationary phase) without concentration of biomass, as the enrichment completely lost ammonia oxidation activity after concentration attempts by centrifugation. In contrast, in the present study, pure active '*Ca.* N. uzonensis' N4 biomass was successfully concentrated using Amplicon filter units (see details in Materials and Methods in the main text). Previously, it has been observed that the substrate affinity of *Nitrobacter winogradsky* differs by more than an order of magnitude between biomass harvested during exponential phase versus stationary phase [26]. In addition, the '*Ca.* N. uzonensis' N4 enrichment culture previously tested in 2017 was re-examined after being cultured at 37°C and harvested using the Amicon filter units. Using these conditions, there was no significant difference ($p$ = 0.95) between the substrate affinity for $NH_3$ observed between the enrichment culture ($K_{m(app)}$=18.05 nM, s.d.=9.55) and the pure culture ($K_{m(app)}$=19.1 nM, s.d.=6.98; Supplementary Fig. 1), but we cannot rule out that the composition of the enrichment culture changed between 2017 and our study.

A second possible explanation for the observed differences between the enrichment culture and the pure culture is an effect of temperature on '*Ca.* N. uzonensis' N4 cellular kinetics, as the measurements were previously performed at 46°C [19], versus 37°C in this study. Indeed, increased substrate affinities at lower temperatures were also observed in the present study for '*Ca*. N. oleophilus' MY3 (Supplementary Fig. 7) and for *N. viennensis* EN76 (~400 nM at 42 °C versus ~20-60 nM $NH_3$ at 30°C) across two independent studies [19,27]. In the latter example, there were no large differences in growth medium or culture pH, but one study was performed at the optimal growth (determined by nitrite production) temperature of 42°C [19], whereas the other study was performed at the more environmentally relevant temperature of 30°C [27]. However, neither of these examples display the variation observed for '*Ca*. N. uzonensis' N4. Therefore, we postulate that due to the differences in cell harvesting techniques, the reported kinetic properties of '*Ca*. N. uzonensis' N4 are likely representative of different growth stages.

**Determining cellular activity kinetics with MR versus nitrite production**

The cellular kinetics of '*Ca*. Nitrosocosmicus franklandus' C13 were determined in this study by two different methods, both MR and short-term activity (measured by $NO_2^-$ production) (Supplementary Figs. 2 and 10). The MR experiments were performed at the optimal growth temperature of 42°C [18], while the short-term activity measurements were conducted at both 37°C and 42°C. Notably, there were significant differences ($p < 0.01$) in the observed $K_{m(app)}$ for $NH_3$ between the two methods. As it does not appear to be caused by different assay temperatures ($p = 0.29$), it is possible this difference stems from the different methods the cells were harvested or reflects that the cells were in different growth phases when the kinetic experiments were performed. For the MR experiments, '*Ca*. Nitrosocosmicus franklandus' C13 culture was taken without concentration once a culture depleted all the substrate. This means that the growth phase of the culture was late-log or early stationary phase. In contrast, for the short-term activity assays, '*Ca*.

Nitrosocosmicus franklandus' C13 cultures were harvested (filtered, see above) when in log phase. While these differences in the $K_{m(app)}$ for $NH_3$ of '*Ca*. Nitrosocosmicus franklandus' C13 are significant, they are all well within the range of the $K_{m(app)}$ for $NH_3$ of the other AOA in the *Nitrososphaeraceae* lineage.

Together, these observations highlight the importance of cultivation conditions and the methods used when determining the cellular kinetics of AOM. They raise an interesting question about whether the kinetic properties determined under more *in-situ*-like conditions or at laboratory determined optimal conditions provides more useful information. These differences will need to be considered in future studies, as most AOM are cultivated at temperatures much higher than what is found in their natural environmental habitat. In addition, these observations also highlight the fact that the whole cell (apparent) kinetic properties reported here and elsewhere have inherent plasticity and are not equivalent to kinetic constants. This is important to remember when predicting which AOM may outcompete others in both laboratory and environmental settings.

### Supplementary Tables and Figures:

**Supplementary Table 1.** Growth medium of the ammonia-oxidizing microorganisms used in this study.

| | **Medium #1[a]** | **Medium #2[b]** | **Medium #3[c]** | **Medium #4[d]** | **Medium #5[e]** |
|---|---|---|---|---|---|
| **Alternative name** | - | Artificial freshwater medium | Synthetic *Crenarchaeote* medium | - | - |
| **Cultures** | ‘*Ca*. N. uzonensis’ N4<br>*N. inopinata*<br>*N. europaea* | ‘*Ca*. N. chungbukensis’ MY2<br>‘*Ca*. N. oleophilus’ MY3<br>‘*Ca*. N. franklandus’ C13<br>*N. koreense* MY1 | *N. maritimus* SCM1<br>*N. piranensis* D3C<br>*N. adriaticus* NF5 | ‘*Ca*. N. devanaterra’ Nd1<br>‘*Ca*. N. sinensis’ Nd2 | ‘*Ca*. N. yellowstonensis’ HL72<br>‘*Ca*. N. tenchongensis’ DRC1<br>‘*Ca*. N. nevadensis’ GerE |
| **Contents (g $l^{-1}$)** | **(g $l^{-1}$)**<br>NaCl (0.6)<br>$MgSO_4 \times 7\ H_2O$ (0.05)<br>$KH_2PO_4$ (0.05)<br>KCl (0.075) | **(g $l^{-1}$)**<br>NaCl (1.0)<br>$MgCl_2 \times 6H_2O$ (0.4)<br>$CaCl_2 \times 2H_2O$ (0.1)<br>$KH_2PO_4$ (0.2)<br>KCl (0.5) | **(g $l^{-1}$)**<br>NaCl (26)<br>$MgSO_4 \times 7H_2O$ (5)<br>$MgCl_2 \times 6H_2O$ (5)<br>$CaCl_2 \times 2H_2O$ (1.5)<br>Kbr. (0.1) | **(g $l^{-1}$)**<br>NaCl (1)<br>$MgCl_2 \times 6H_2O$ (0.4)<br>$CaCl_2 \times 2H_2O$ (0.1)<br>$KH_2PO_4$ (0.2)<br>KCl (0.5) | **(g $l^{-1}$)**<br>NaCl (1)<br>$MgCl_2 \times 6H_2O$ (0.4)<br>$CaCl_2 \times 2H_2O$ (0.1)<br>KCl (0.5) |
| **Additional contents (stock solutions)** | **Trace element solution contents (TES) (mg $l^{-1}$):**<br>HCl (37%) (2.5 ml $l^{-1}$)<br>$MnSO_4 \times 1\ H_2O$ (34.4)<br>$H_3BO_3$ (50.0)<br>$ZnCl_2$ (70.0)<br>$Na_2MoO_4 \times 2\ H_2O$(72.6)<br>$CuCl_2 \times 2\ H_2O$ (20.0)<br>$NiCl_2 \times 6\ H_2O$ (24.0)<br>$CoCl_2 \times 6\ H_2O$ (80.0)<br>$FeSO_4 \times 7\ H_2O$ (1000)<br>**Selenium wolfram solution (SWS) (mg $l^{-1}$):**<br>$Na_2SeO_3 \times 5H_2O$ (3.0)<br>$Na_2WO_4 \times 2H_2O$ (4.0)<br>NaOH. (500)<br>**$CaCl_2$ solution (g $l^{-1}$):**<br>CaCl2 x 2H2O (147) | **FeNaEDTA (2.75 g $l^{-1}$)**<br>**Non-chelated TES (mg $l^{-1}$):**<br>HCI (25%) (12.5 ml $l^{-1}$)<br>$FeSO_4 \times 7H20$ (2100)<br>$H_3BO_3$ (30)<br>$MnC1_2 \times 4H_20$ (100)<br>$CoC1_2 \times 6H_20$ (190)<br>$NiC1_2 \times 6H_20$ (24)<br>$CuC1_2 \times 2H_20$ (2)<br>$ZnSO_4 \times 7H_20$ (144)<br>$Na_2MoO_4 \times 2H20$ (36)<br>**Vitamin solution (mg $l^{-1}$):**<br>Biotin (20)<br>Folic Acid (20)<br>Pyridoxine HCl (100)<br>Thiamine HCl (50)<br>Riboflavin (50)<br>Nicotinic Acid (50)<br>DL Pantothenic Acid (50)<br>p Aminobenzoic Acid (50)<br>Lipoic acid (50)<br>1,4-naphthaquinine (40) | **TES (see Medium #1)**<br>**SWS (see Medium #1)**<br>**HEPES (238 g $l^{-1}$)**<br>**$NaHCO_3$ (84 g $l^{-1}$)**<br>**$KH_2PO_4$ (0.4 g $l^{-1}$)**<br>**FeNaEDTA (2.75 g $l^{-1}$)**<br>**Na-pyruvate (110 g $l^{-1}$)** | **TES (see Medium #1)**<br>**FeNaEDTA (2.75 g $l^{-1}$)**<br>**MES hydrate**<br>**Oxalacetic acid (66 g $l^{-1}$)** | **TES (see Medium #1)**<br>**SWS (see Medium #1)**<br>**$NaHCO_3$ (84 g $l^{-1}$)**<br>**MOPS (209.2 g $l^{-1}$) pH7.5**<br>**$KH_2PO_4$ (136 g $l^{-1}$)**<br>**FeNaEDTA (2.75 g $l^{-1}$)**<br>**Vitamin solution (mg $l^{-1}$):**<br>Biotin (20)<br>Folic Acid (20)<br>Pyridoxine HCl (100)<br>Thiamine HCl (50)<br>Riboflavin (50)<br>Nicotinic Acid (50)<br>DL Pantothenic Acid (50)<br>p Aminobenzoic Acid (50)<br>Choline Chloride (2000)<br>Vitamin $B_{12}$ (10)<br>pH =7 with KOH |

| | | | | | |
|---|---|---|---|---|---|
| | | Nicotinamide (100)<br>Hemin (10)<br>Vitamin $B_{12}$ (10)<br>pH =7 with KOH | | | |
| **Procedure** | Contents were autoclaved and the additional contents were aseptically added afterwards (ml $l^{-1}$):<br>• TES (1)<br>• SWS. (1)<br>• $CaCl_2$ (1) | Contents were autoclaved and the additional contents were added aseptically afterwards (ml $l^{-1}$):<br>• Non-chelated TES (1)<br>• Vitamin solution. (1)<br>• FeNaEDTA. (1)<br>• HEPES (10)<br>• $NaHCO_3$ (2) | Contents were autoclaved and the additional contents were added aseptically afterwards (ml $l^{-1}$):<br>• HEPES (10)<br>• $NaHCO_3$ (2)<br>• $KH_2PO_4$ (5)<br>• FeNaEDTA (1)<br>• TES (1)<br>• Na-pyruvate (1) | Contents and additional contents were combined and then filter sterilized (0.22µm). The amount of additional contents added were:<br>• Oxalacetate (1 ml $l^{-1}$)<br>• FeNaEDTA (1 ml $l^{-1}$)<br>• TES (1 ml $l^{-1}$)<br>• MES hydrate (1.95 g) | Contents were autoclaved and the additional contents were added aseptically afterwards (ml $l^{-1}$):<br>• TES (10)<br>• SWS (1)<br>• $NaHCO_3$ (5)<br>• $KH_2PO_4$ (0.3)<br>• FeNaEDTA (1)<br>• Vitamin solution (1)<br>• MOPS. (10) |

[a]Medium was modified from Kits *et al*., 2017 [19]
[b]Medium was modified from Jung *et al*., 2011 [20]
[c]Medium was modified from Könnecke *et al*., 2005 [28]
[d]Medium was modified from Lehtovirta-Morley *et al*., 2011 [29]
[e]Medium was modified from de la Torre *et al*., 2008 [12]

**Supplementary Table S2**. Optimal growth conditions and the conditions used for cellular kinetic experiments in this study for all tested ammonia-oxidizing microorganisms. References are provided for optimal growth conditions. Cellular kinetics were determined at the temperature and pH ranges noted in parenthesis, if different than optimal growth conditions.

| Type of nitrifier | | | | Growth optimum | | | | |
|---|---|---|---|---|---|---|---|---|
| **Type** | **Group** | **Culture** | | **pH** | **Temperature (℃)** | **Medium[a]** | **Isolation source** | **Reference[b]** |
| AOA | *Nitrosopumilales* | *Nitrosopumilus maritimus* SCM1 | Pure | 7.3 (8) | 32 (30) | #3 | Marine aquarium | 28,30 |
| | | *Nitrosopumilus piranensis* D3C | Pure | 7.3 (7) | 30 (30) | #3 | Marine water | 31 |
| | | *Nitrosopumilus adriaticus* NF5 | Pure | 7.3 (7) | 30 (30) | #3 | Marine water | 31 |
| | | *Nitrosarchaeum koreense* MY1 | Pure | 7.0 (7.2) | 25 (25) | #2 | Agricultural soil | 32 |
| | | '*Ca.* Nitrosotenuis chungbukensis' MY2 | Pure | 7.0 (7.3) | 30 (30) | #2 | Agricultural soil | 33 |
| | | '*Ca.* Nitrosotenuis uzonensis' N4[c] | Pure | 7.5 (7.1-7.4) | 37 (37) | #1 | Thermal spring | 1, this study |
| | | | Enrichment | (7.75) | (37) | | | |
| | '*Ca.* Nitrosotaleales' | '*Ca.* Nitrosotalea devanaterra' Nd1 | Pure | 5.3 | 30 | #4 | Acidic soil | 34 |
| | | '*Ca.* Nitrosotalea sinensis' Nd2 | Pure | 5.3 | 35 | #4 | Acidic soil | 34 |
| | *Nitrososphaerales* | *Nitrososphaera viennensis* EN76 | Pure | 7.6 | 42 | -[d] | Garden soil | 35,36 |
| | | '*Ca.* Nitrososphaera gargensis' | Pure | 7.5 | 42 | -[d] | Thermal spring | 37 |
| | | '*Ca.* Nitrosocosmicus oleophilus' MY3 | Pure | 7.0 (6.5-8.3) | 30 (30) | #2 | Oil contaminated sediment | 38 |
| | | '*Ca.* Nitrosocosmicus franklandus' C13 | Pure | 7.8 (7.1-8) | 42 (37-42) | #2 | Agricultural soil | 18 |
| | | '*Ca.* Nitrososphaera nevadensis' GerE[3] | Enrichment | 7.1 (7.35-7.5) | 50 (50) | #5 | Thermal spring | This study |
| | '*Ca.* Nitrosocaldales' | '*Ca.* Nitrosothermus tengchongensis' DRC1[3] | Enrichment | 7.1 (7.3-7.5) | 72 (72) | #5 | Thermal spring | This study |
| | | '*Ca.* Nitrosocaldus yellowstonensis' HL72[3] | Enrichment | 7.1 (7.3-7.5) | 72 (72) | #5 | Thermal spring | 12 |
| Comammox | Clade A | *Nitrospira inopinata* | Pure | 7.5 (6.7-8.4) | 37 (37) | #1 | Thermal spring | 19 |

| | | | | | | | | |
|---|---|---|---|---|---|---|---|---|
| AOB | Beta-proteobacteria | *Nitrosomonas europaea* ATCC19718 | Pure | 7.5 | 25 | #1 | Soil | 20,26 |

[a]Medium contents are described in Supplementary Table 1

[b]References refer to the physiological characterization of optimum growth pH and temperature for each AOM strain

[c]The pH and temperature optimum have not been previously reported for these AOM strains

[d]*N. gargensis* and *N. viennensis* were not analyzed in this study, but are included here for comparison

**Supplemental Table S3.** The effect of medium pH and temperature on the cellular kinetic properties of selected ammonia-oxidizing microorganisms.

| Culture | Temperature (°C) | Cell preparation | pH | Number of replicates | Average measured $K_{m(app)}$ for $NH_3 + NH_4^+$ (μM) | Average calculated $K_{m(app)}$ for $NH_3$ (nM) | $V_{max}$ (μmol N mg protein$^{-1}$ h$^{-1}$)[a] | Reference |
|---|---|---|---|---|---|---|---|---|
| '*Ca.* Nitrosocosmicus oleophilus' MY3[b] | 30 | Whole-cell | 6.5 | 6 | 2516.87 ± 647.78 | 6630.11 ± 935.50 | 37.04 ± 6.41 | This study |
| | | | 7.2 | 6 | 818.73 ± 185.44 | 9792.16 ± 1322.34 | 36.12 ± 5.13 | |
| | | | 8.3 | 5 | 246.29 ± 97.82 | 26284.80 ± 3460.81 | 35.84 ± 5.01 | |
| | 25 | | 7.2 | 2 | 879.23 ± 10.98 | 7953.70 ± 98.11 | 21.97 ± 1.32 | |
| | 30 | | | 2 | 970 ± 19.42 | 12233 ± 244.63 | 35.11 ± 0.11 | |
| | 35 | | | 2 | 832.42 ± 15.01 | 14607.28 ± 264.41 | 47.75 ± 2.88 | |
| *Nitrospira inopinata*[b] | 37 | Whole-cell | 6.7 | 6 | 2.24 ± 0.53 | 12.32 ± 1.55 | 12.35 ± 0.46 | This study |
| | | | 7.1 | 4 | 0.97 ± 0.17 | 15.47 ± 2.69 | 14.77 ± 1.35 | |
| | | | 7.8 | 4 | 0.39 ± 0.09 | 30.33 ± 7.61 | 10.42 ± 2.09 | |
| | | | 8.4 | 4 | 0.24 ± 0.08 | 36.99 ± 12.12 | 10.56 ± 0.85 | |
| *Nitrosomonas europaea* | 25 | Whole-cell | 7.0 | 1 | 4000 | 2300 | N/A | 39 |
| | | | 7.5 | 1 | 1600 | 2900 | N/A | |
| | | | 8.0 | 1 | 480 | 2600 | N/A | |
| | | | 8.5 | 1 | 300 | 4600 | N/A | |
| | | | 9.1 | 1 | 140 | 5800 | N/A | |
| | | Cell-free extract | 6.5 | 1 | 10000 | 18000 | N/A | |
| | | | 7.0 | 1 | 4000 | 23000 | N/A | |
| | | | 7.5 | 1 | 1300 | 24000 | N/A | |

| | | | | | |
|---|---|---|---|---|---|
| | 8.0 | 1 | 320 | 18000 | N/A |
| | 8.5 | 1 | 120 | 20000 | N/A |

[a]$V_{max}$ data unavailable for experiment is denoted (N/A)
[b]The individual replicates are plotted in Fig. 4 and supplementary Fig. 7

**Supplementary Table S4.** Cellular morphology of ammonia oxidizers. All cell diameter and lengths were obtained from previously published images. TEM and SEM cell sizes were taken directly from the references cited. MicrobeTracker[40] was used to calculate cell sizes when phase contrast images were provided.

| Strain | Shape[a] | Diameter (µm) | Length (µm) | Surface area / volume ratio | Type of images[b] | Reference |
|---|---|---|---|---|---|---|
| ***Nitrosopumilales* (Group I.1a)** | | | | | | |
| *Nitrosopumilus maritimus* SCM1 | Rod | 0.17-0.22 | 0.5-0.9 | 13.11 ± 1.83 | TEM, SEM | 28,30 |
| *Nitrosopumilus piranensis* D3C | Rod | 0.2-0.25 | 0.49-2.0 | 11.44 ± 2.04 | TEM, SEM | 31,41 |
| *Nitrosopumilus adriaticus* NF5 | Rod | 0.2-0.25 | 0.59-1.74 | 10.85 ± 1.52 | TEM, SEM | 31,41 |
| *Nitrosacheum koreensis* MY1 | Rod | 0.3-0.6 | 0.6-1.0 | 9.05 ± 2.47 | TEM, SEM | 20,32 |
| ‘*Ca.* Nitrosotenuis chungbukensis’ MY2 | Rod | 0.2-0.35 | 0.6-1.2 | 7.71 ± 2.73 | TEM, SEM | 33 |
| ‘*Ca.* Nitrosotenuis uzonensis’ N4 | Rod | 0.2-0.3 | 0.5-1.7 | 8.10 ± 2.84 | TEM | 1 |
| **‘*Ca.* Nitrosotaleales’ (Group I.1a-associated)** | | | | | | |
| ‘*Ca.* Nitrosotalea devanaterra’ Nd1 | Rod | 0.28-0.33 | 0.5-1 | 9.80 ± 1.26 | TEM, SEM | 29,34 |
| ‘*Ca.* Nitrosotalea sinensis’ Nd2 | Rod | 0.23-0.28 | 0.6-0.85 | 9.61 ± 1.17 | TEM, SEM | 34 |
| ***Nitrososphaerales* (Group I.1b)** | | | | | | |
| ‘*Ca.* Nitrososphaera gargensis’[c] | Coccus | 0.45 | 0.45 | 6.66 | TEM | 42 |
| *Nitrososphaera viennensis* EN76 | Coccus | 0.6-0.8 | 0.6-0.8 | 6.05 ± 0.31 | TEM, SEM, Phase contrast | 35,36 |
| ‘*Ca.* Nitrosocosmicus oleophilus’ MY3 | Coccus | 1.0-1.1 | 1.0-1.1 | 3 ± 0.15 | TEM, SEM | 38 |
| ‘*Ca.* Nitrosocosmicus franklandus’ C13 | Coccus | 0.85-1.2 | 0.85-1.2 | 2.72 ± 0.44 | TEM, SEM | 18 |
| **Ammonia-oxidizing bacteria** | | | | | | |
| *Nitrosococcus oceani* | Coccus | 1.3-1.5 | 1.4-1.5 | 2.07 ± 0.06 | TEM | 43 |
| *Nitrosomonas europaea* | Rod | 0.8-1.1 | 1.0-1.7 | 2.99 ± 0.85 | TEM, Phase contrast | 44,45 |
| *Nitrosomonas oligotropha* | Rod | 0.5-0.7 | 1.1-2.1 | 5.18 ± 0.72 | Phase contrast | 45 |
| *Nitrosomonas eutropha* | Rod | 0.85-1.2 | 2.6-2.9 | 2.38 ± 0.17 | Phase contrast | 45 |
| *Nitrosospira briensis* | Spiral | 0.3-0.45 | 1.0-1.2 | 6.82 ± 0.37 | TEM, SEM | 44,46 |
| '*Ca.* Nitrosoglobus terrae' | Coccus | 2-3 | 2-3 | 1.2 ± 0.21 | TEM, Phase contrast | 47 |
| ‘*Ca.*Nitrosacidococcus tergens’[3] | Coccus | 0.5 | 0.5 | 6 | TEM | 48 |
| *Nitrosomonas cryotolerans* | Rod | 1.2-2.2 | 2-4 | 1.84 ± 0.52 | TEM | 49 |
| *Nitrosomonas communis* | Rod | 1-1.4 | 1.7-2.2 | 2.69 ± 0.34 | Phase contrast | 45 |
| **Comammox bacteria** | | | | | | |
| *Nitrospira inopinata* | Spiral | 0.18-0.3 | 0.6-1.7 | 10.15 ± 2.74 | TEM, SEM | 19,50 |

[a]The spiral shaped *N. briensis* and ‘*Ca.* N. inopinata’ were treated as rods
[b]TEM, transmission electron microscopy; SEM, scanning electron microscopy
[c]The length and diameter of ‘*Ca.* N. gargensis’ and ‘*Ca.* N. tergens’ are each representative of a single cell measurement

**Supplementary Fig. S1. Ammonia oxidation kinetics of *Nitrosopumilales* (Group I.1a) AOA.** Michaelis-Menten plots of *N. maritimus* SCM1, *N. piranensis* D3C, *N. adriaticus* NF5, *N. koreense* MY1, '*Ca*. N. chungbukensis' MY2, and '*Ca*. N. uzonensis' N4. Total ammonium oxidation rates were determined from microsensor measurements of substrate dependent $O_2$ consumption from either discrete slopes over many substrate concentrations (a,e,f,h,i,k,o,p,r,v) or a single trace measurement (b,c,d,g,j,l,m,n,q,s,t,u). Apparent half-saturation ($K_{m(app)}$) and maximum oxidation rates ($V_{max}$) for total ammonium were calculated by fitting the data to the Michaelis-Menten kinetic equation. The red line indicates the best fit of the data. Standard deviations of the estimates based on the non-linear regression are reported. Microrespiration conditions for each strain are presented in Supplementary Table 2.

***Nitrosopumilus maritimus* SCM1**

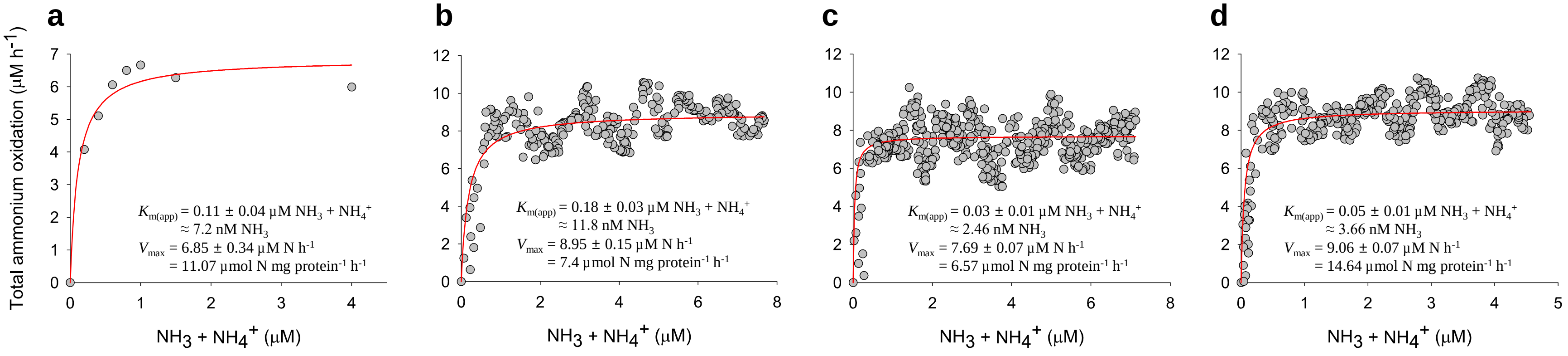


***Nitrosopumilus piranensis* D3C**

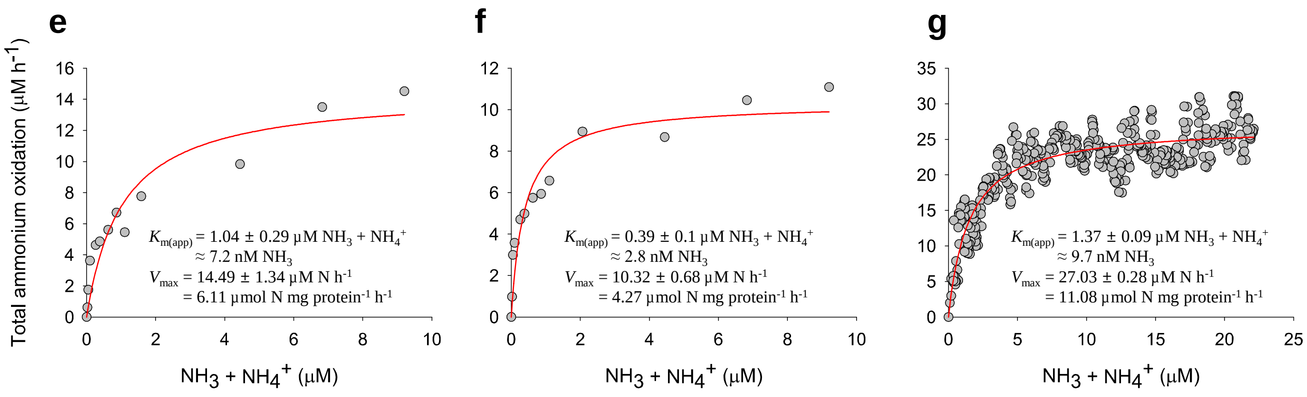

### *Nitrosopumilus adriaticus* NF5

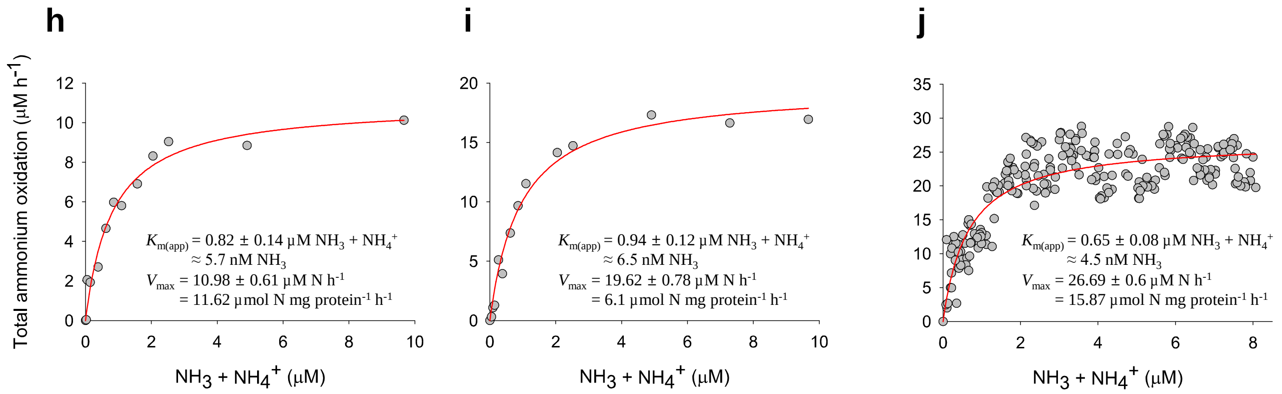


### *Nitrosopumilus koreense* MY1

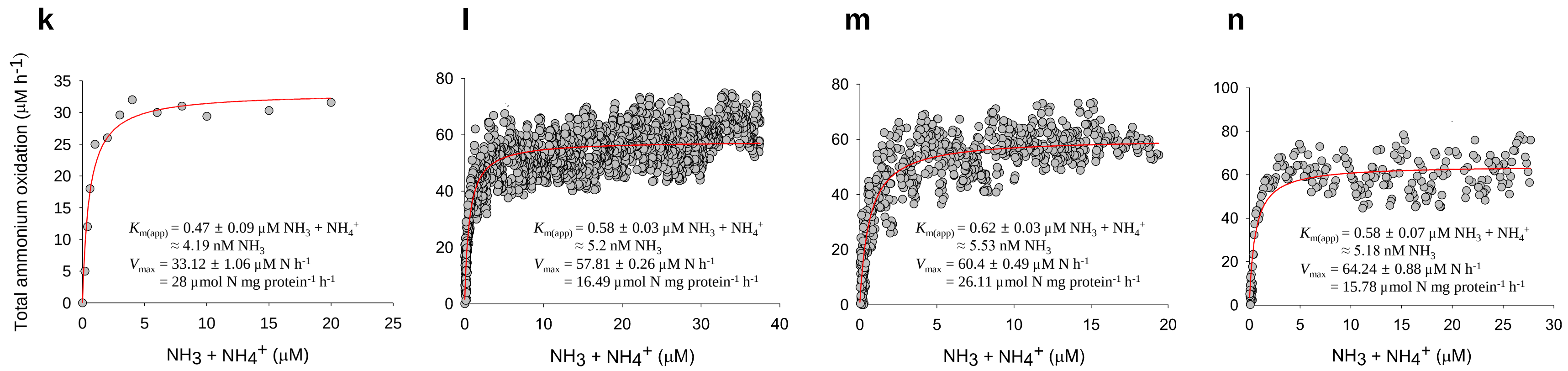

### ‘*Ca*. Nitrosotenuis chungbukensis’ MY2

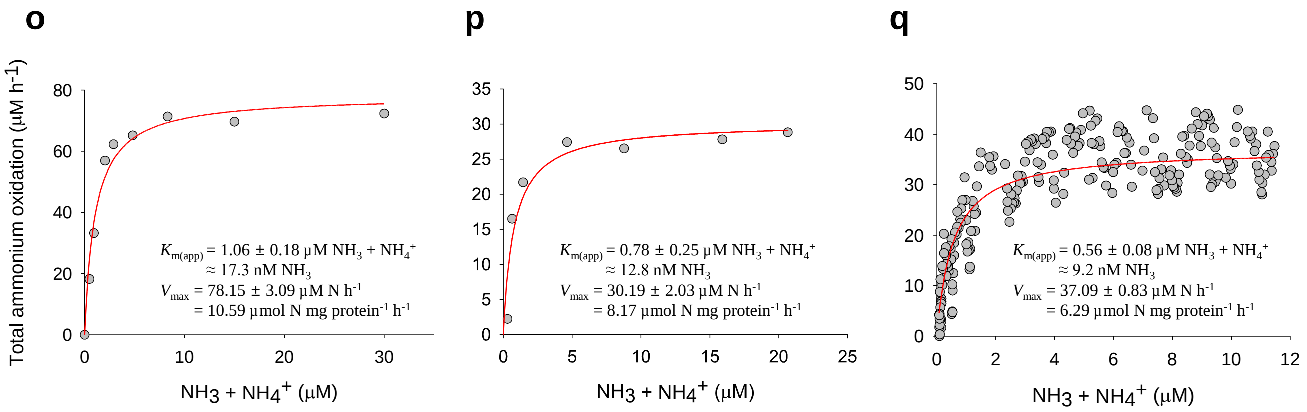


### ‘*Ca.* Nitrosotenuis uzonensis’ N4 (pure culture)

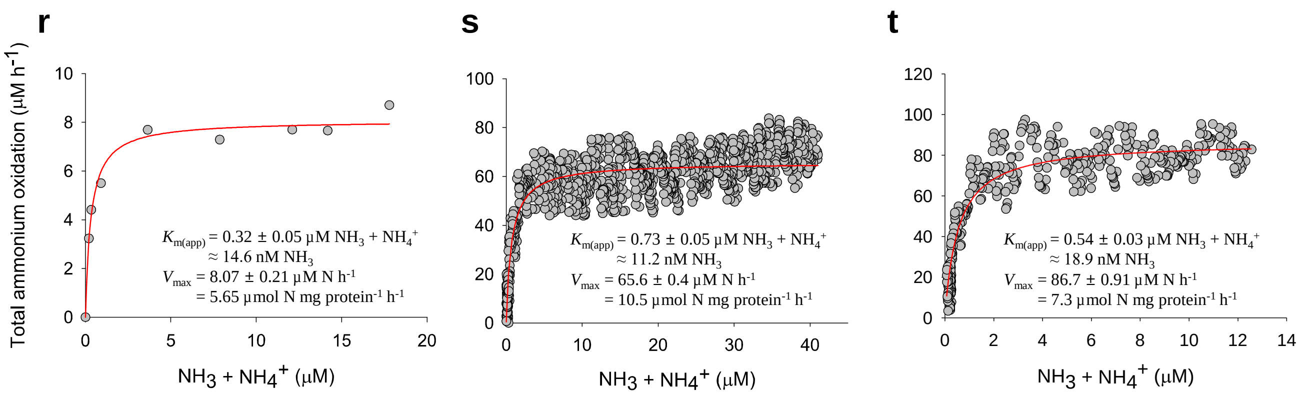

**'*Ca.* Nitrosotenuis uzonensis' N4 (enrichment culture)**

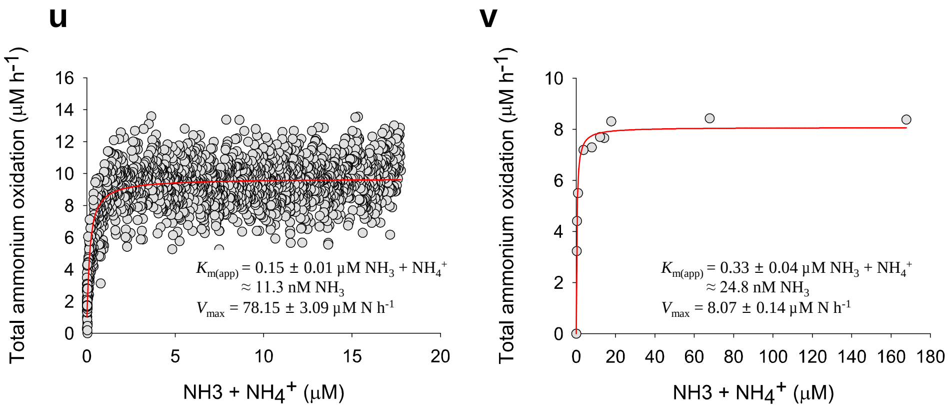

**Supplementary Fig. S2. Comparison of AOA apparent substrate affinity ($K_{m(app)}$) values across studies.** The (a) $K_{m(app)}$ for ($NH_3$) and (b) $K_{m(app)}$ for total ammonium, of *N. maritimus* SCM1 (brown), *N. koreensis* MY1 (orange), '*Ca*. N. uzonensis' N4 (blue), *N. viennensis* EN76 (purple), *N. inopinata* (green), and '*Ca*. N. franklandus' C13 (yellow) are shown. Symbols filled with light grey represent published values from previous studies and the reference for each strain are indicated. $K_{m(app)}$ values were derived from measurements with either pure (circles) or enrichment (diamonds) cultures. Significant differences between the different experiment for each strain was determined by a Student's t-test: *$p < 0.05$, **$p < 0.005$, *** $p < 0.0005$.

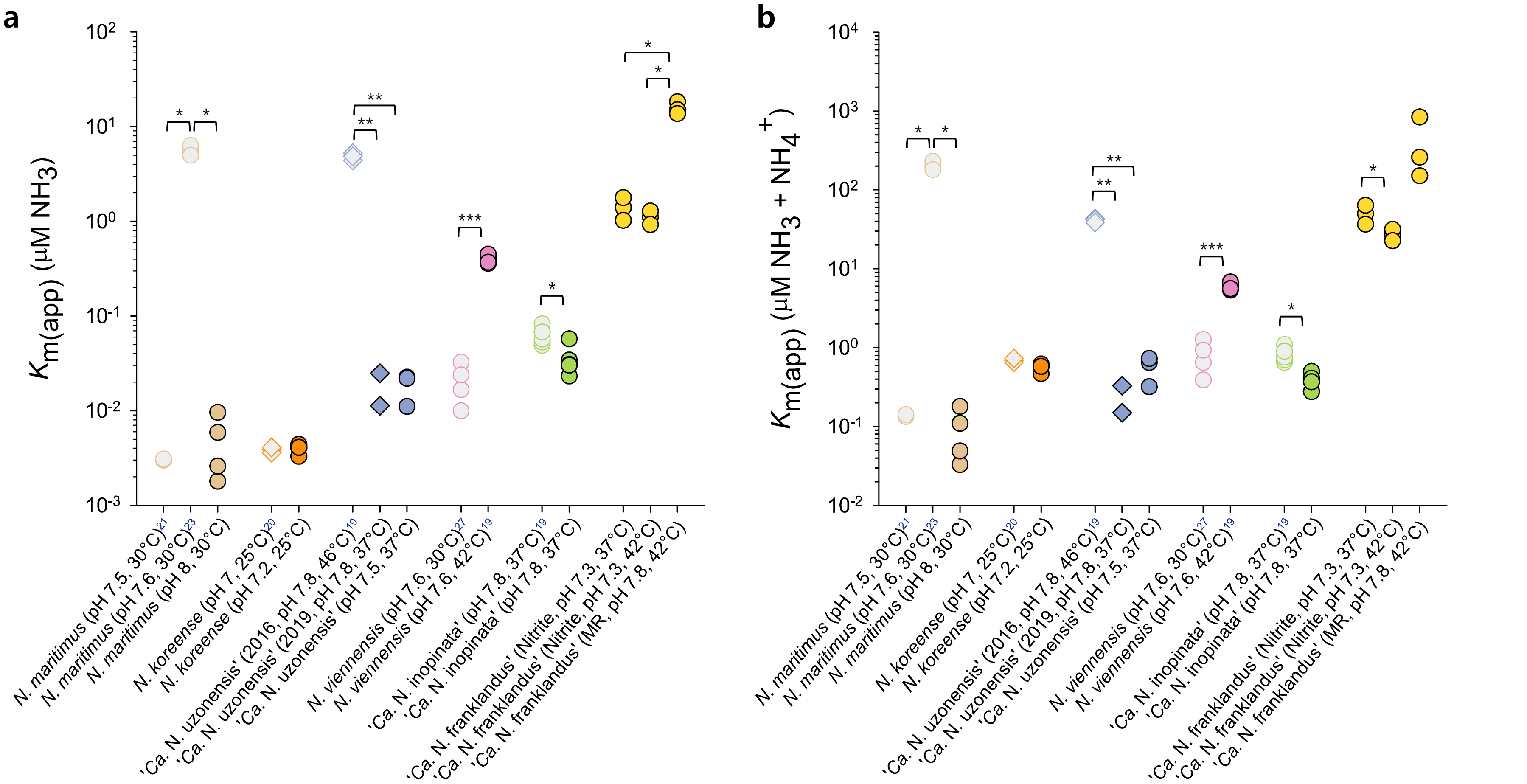

**Supplementary Fig. S3. Ammonia oxidation kinetics of '*Ca.* Nitrosotaleales' (Group I.1a-associated) AOA.** Michaelis-Menten plots for '*Ca*. N. devanaterra' Nd1 and '*Ca*. N. sinensis' Nd2. Total ammonium oxidation rates were determined from microsensor measurements of substrate dependent $O_2$ consumption from discrete slopes over many substrate concentrations. Apparent half-saturation ($K_{m(app)}$) and maximum oxidation rates ($V_{max}$) for total ammonium were calculated by fitting the data to the Michaelis-Menten kinetic equation. The red line indicates the best fit of the data. Standard deviations of the estimates based on the non-linear regression are reported. Microrespiration conditions for each strain are presented in Supplementary Table 2.

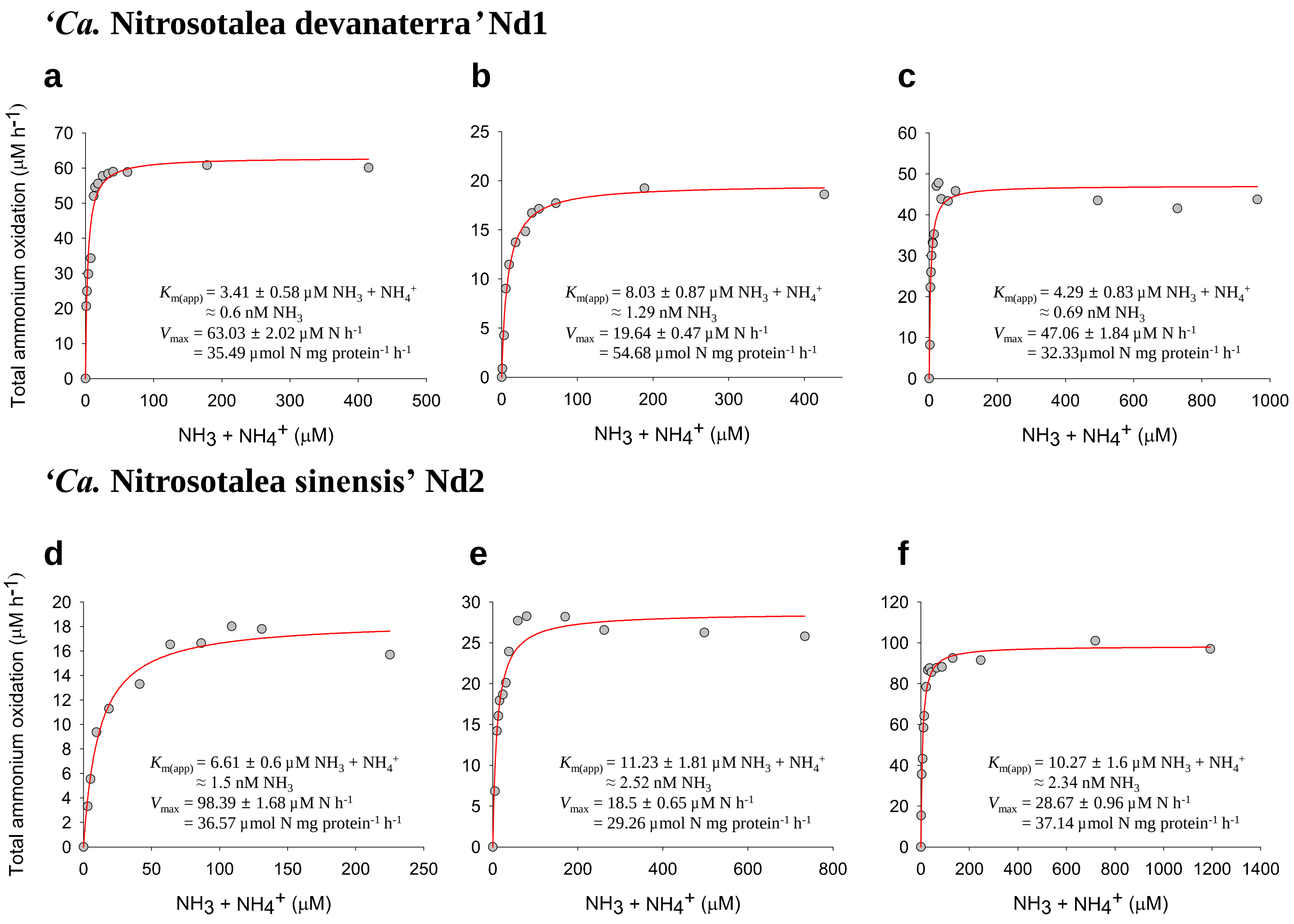

**Supplementary Fig. S4. Ammonia oxidation kinetics of *Nitrososphaerales* (Group I.1b) AOA.** Michaelis-Menten plots for '*Ca*. N. nevadensis' GerE, '*Ca*. N. oleophilus' MY3, and '*Ca*. N. franklandus' C13. Total ammonium oxidation rates were determined from microsensor measurements of substrate dependent $O_2$ consumption from either single trace measurements (a-c) or discrete slopes over many substrate concentrations (d-l). Apparent half-saturation ($K_{m(app)}$) and maximum oxidation rates ($V_{max}$) for total ammonium were calculated by fitting the data to the Michaelis-Menten kinetic equation. The red line indicates the best fit of the data. Standard deviations of the estimates based on the non-linear regression are reported. Microrespiration conditions for each strain are presented in Supplementary Table 2.

**'*Ca.* Nitrososphaera nevadensis' GerE**

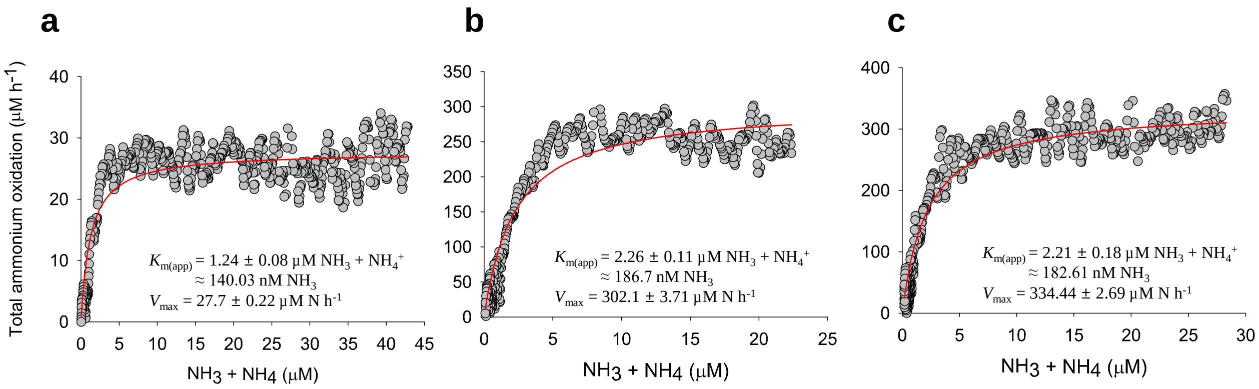


**'*Ca.* Nitrosocosmicus oleophilus' MY3**

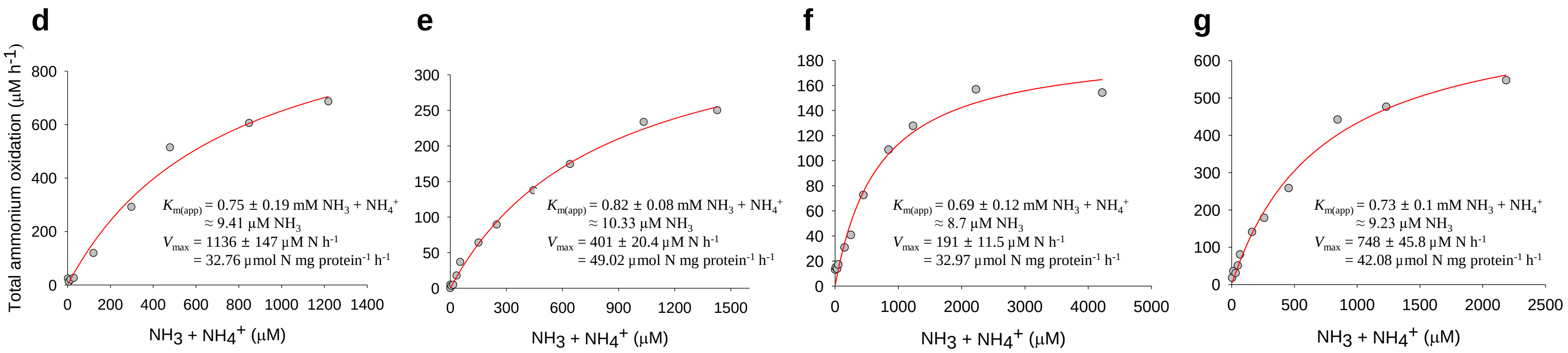

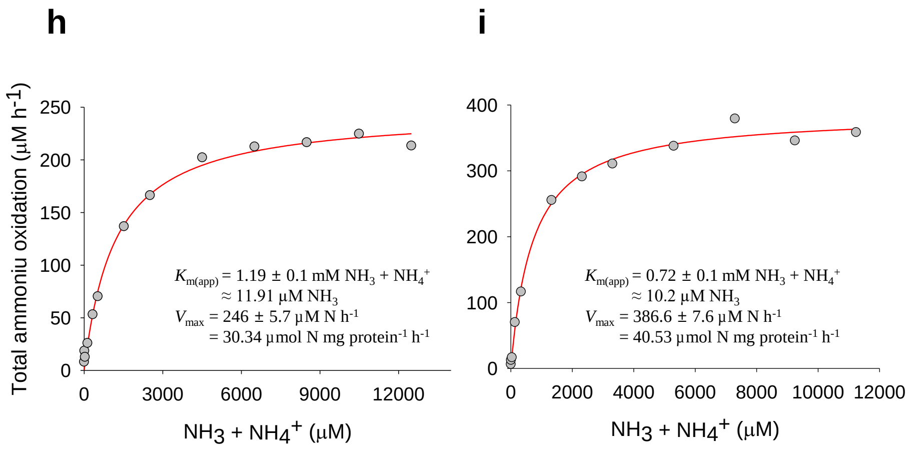


### ‘*Ca.* Nitrosocosmicus franklandus’ C13

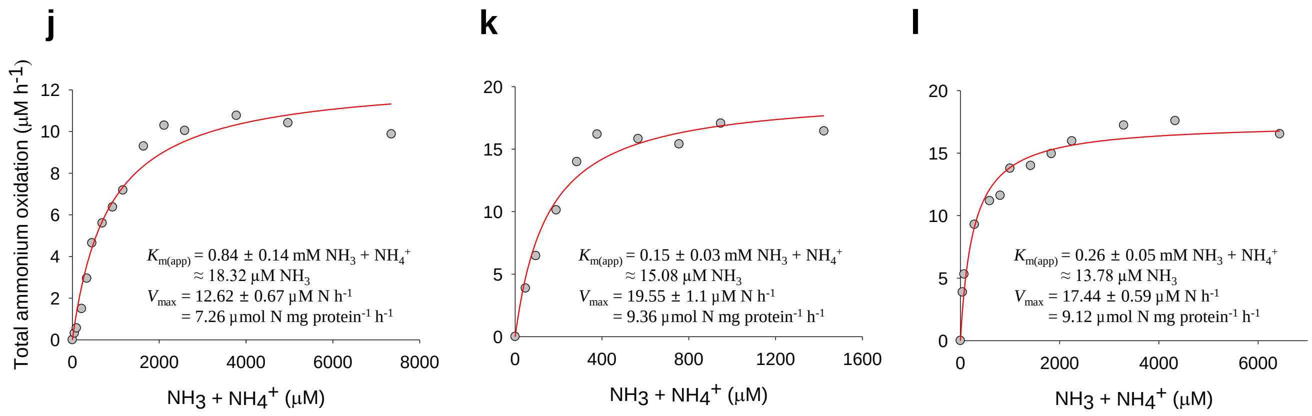

**Supplementary Fig. S5. Ammonia oxidation kinetics of '*Ca.* Nitrosocaldales' (thermophilic) AOA.** Michaelis-Menten plots for '*Ca*. N. tenchongensis DRC1' and '*Ca*. N. yellowstonensis' HL72. Total ammonium oxidation rates were determined from microsensor measurements of substrate dependent $O_2$ consumption from discrete slopes over many substrate concentrations. Only the discrete slopes determined with non-inhibitory ammonium concentrations (highlighted with a red box in each panel) were used to calculate kinetic properties. The apparent half-saturation ($K_{m(app)}$) and maximum oxidation rates ($V_{max}$) for total ammonium were calculated by fitting the data to the Michaelis-Menten kinetic equation. The red line indicates the best fit of the data. Standard deviations of the estimates based on the non-linear regression are reported. Microrespiration conditions for each strain are presented in Supplementary Table 2.

**'*Ca.* Nitrosofervidus tenchongensis' DRC1**

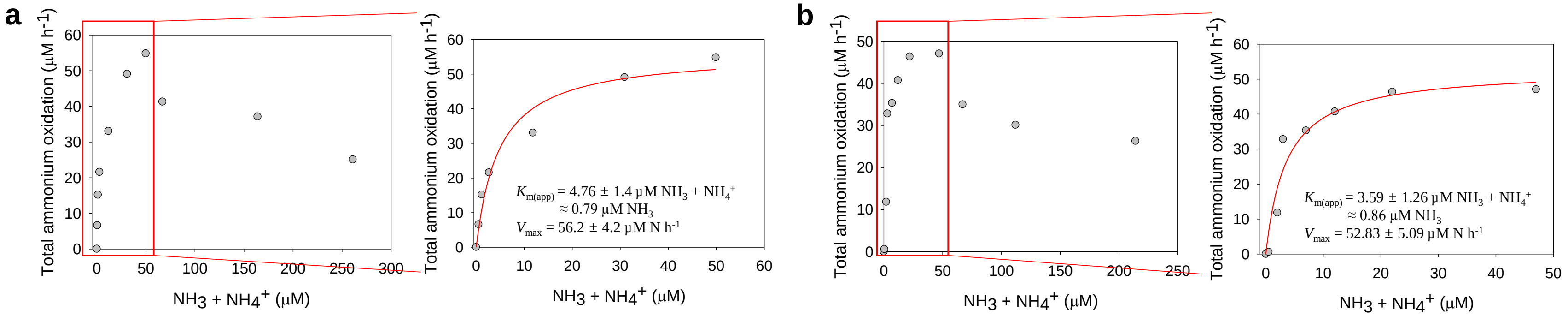


**'*Ca.* Nitrosocaldus yellowstonensis' HL72**

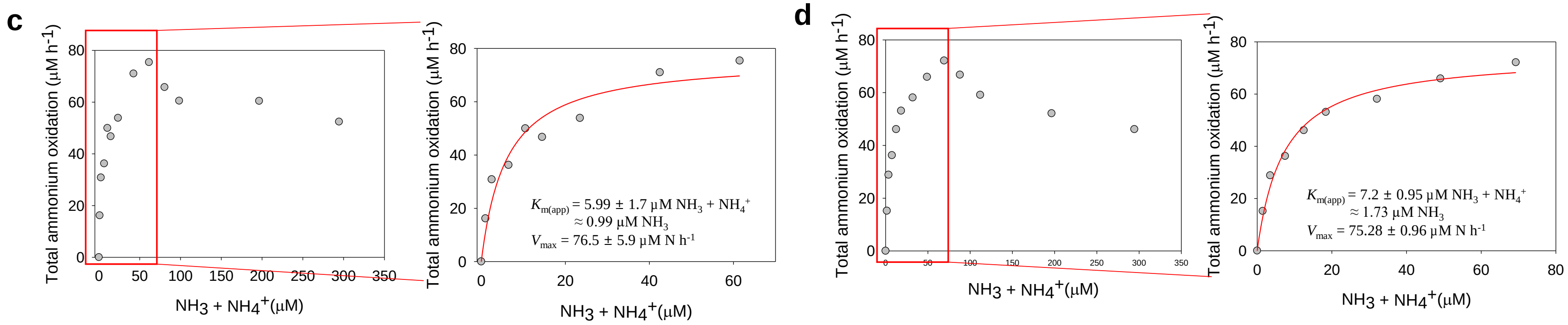

**Supplementary Fig. S6. Effect of short-term temperature shifts on the cellular kinetic properties of '*Ca*. N. oleophilus' MY3.** The (a) apparent substrate affinity ($K_{m(app)}$) for total ammonium, (b) $K_{m(app)}$ for $NH_3$, (c) the specific substrate affinity ($a^o$) for total ammonium, (d) the $a^o$ for $NH_3$ and (e) maximum oxidation rate ($V_{max}$) determined across a range of temperatures is shown. Duplicate experiments are indicated with black and white circles. All experiments were carried out under identical conditions with a constant pH of 7.3. The average $K_{m(app)}$ and $V_{max}$ values for '*Ca*. N. oleophilus MY3' at each temperature are provided in Supplementary Table 3.

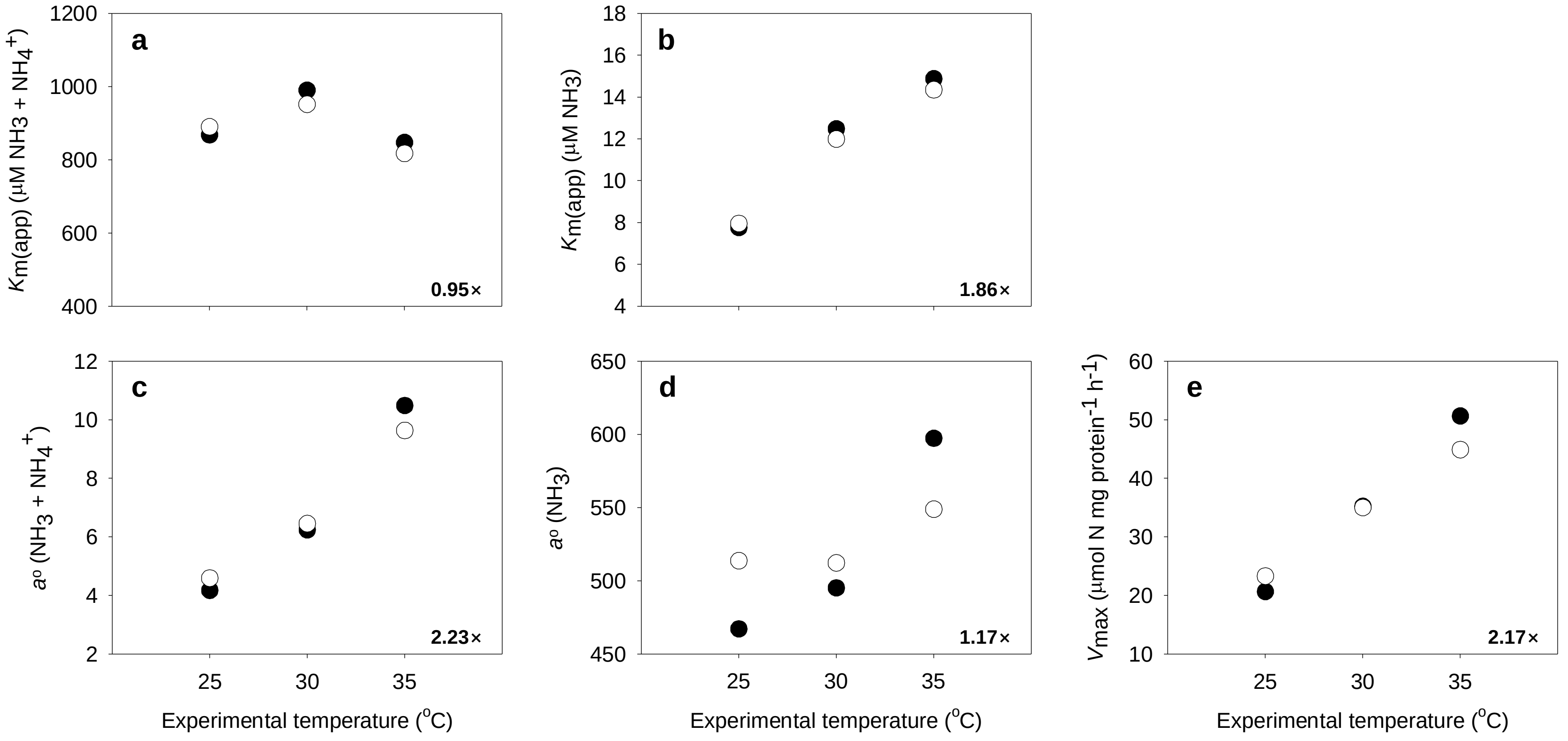

**Supplementary Fig. S7. Microscopic observation of the cell aggregate size of various AOA before (left) and after (right) microrespiration experiments.** DAPI stained (a,b) '*Ca.* Nitrosotenuis uzonensis' N4, (c, d) '*Ca.* N. oleophilus' MY3, and (e, f) *N. piranensis* D3C. Cell aliquots were stained both before (a,c,e) and after (b,d,f) microrespiration experiments. Scale bar 10 µm.

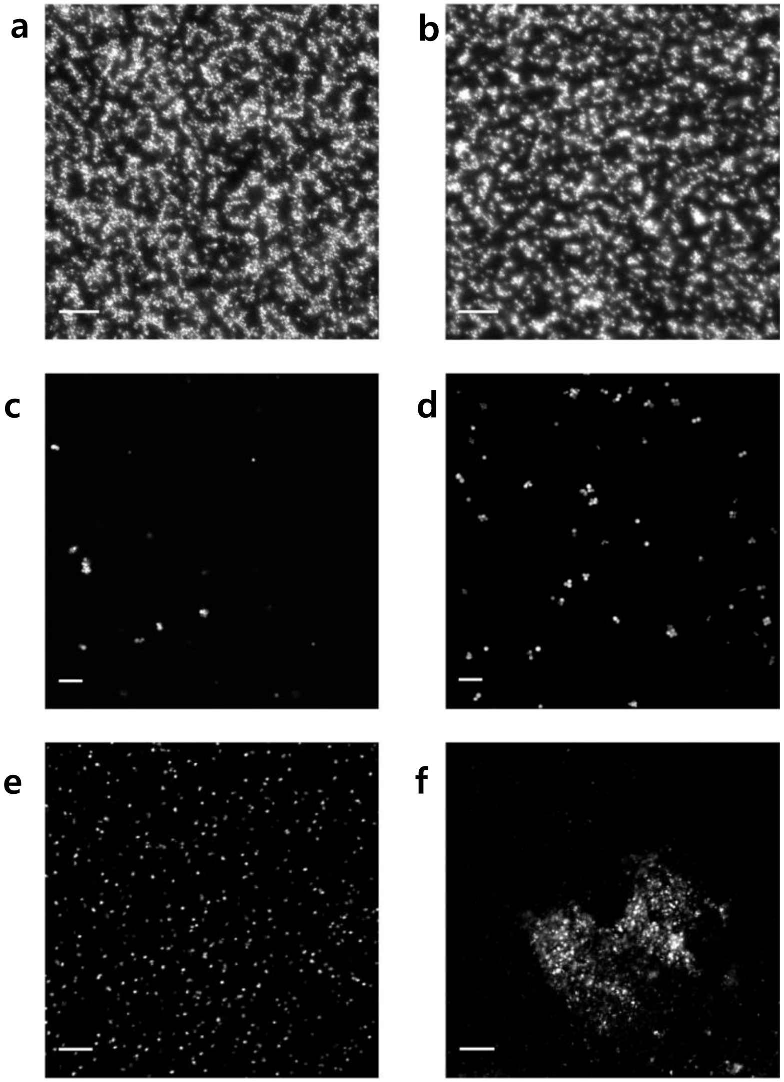

**Supplementary Fig. S8. Ammonia oxidation kinetics of *N. inopinata* and *N. europaea*.** Michaelis-Menten plots for *N. inopinata* (a-d) and *N. europaea* (e,f). Total ammonium oxidation rates were determined from microsensor measurements of substrate dependent $O_2$ consumption from either discrete slopes over many substrate concentrations (a,e,f) or a single trace measurement (b-d). Apparent half-saturation ($K_{m(app)}$) and maximum oxidation rates ($V_{max}$) for total ammonium were calculated by fitting the data to the Michaelis-Menten kinetic equation. The red line indicates the best fit of the data. Standard deviations of the estimates based on the non-linear regression are reported. Microrespiration conditions for each strain are presented in Supplementary Table 2.

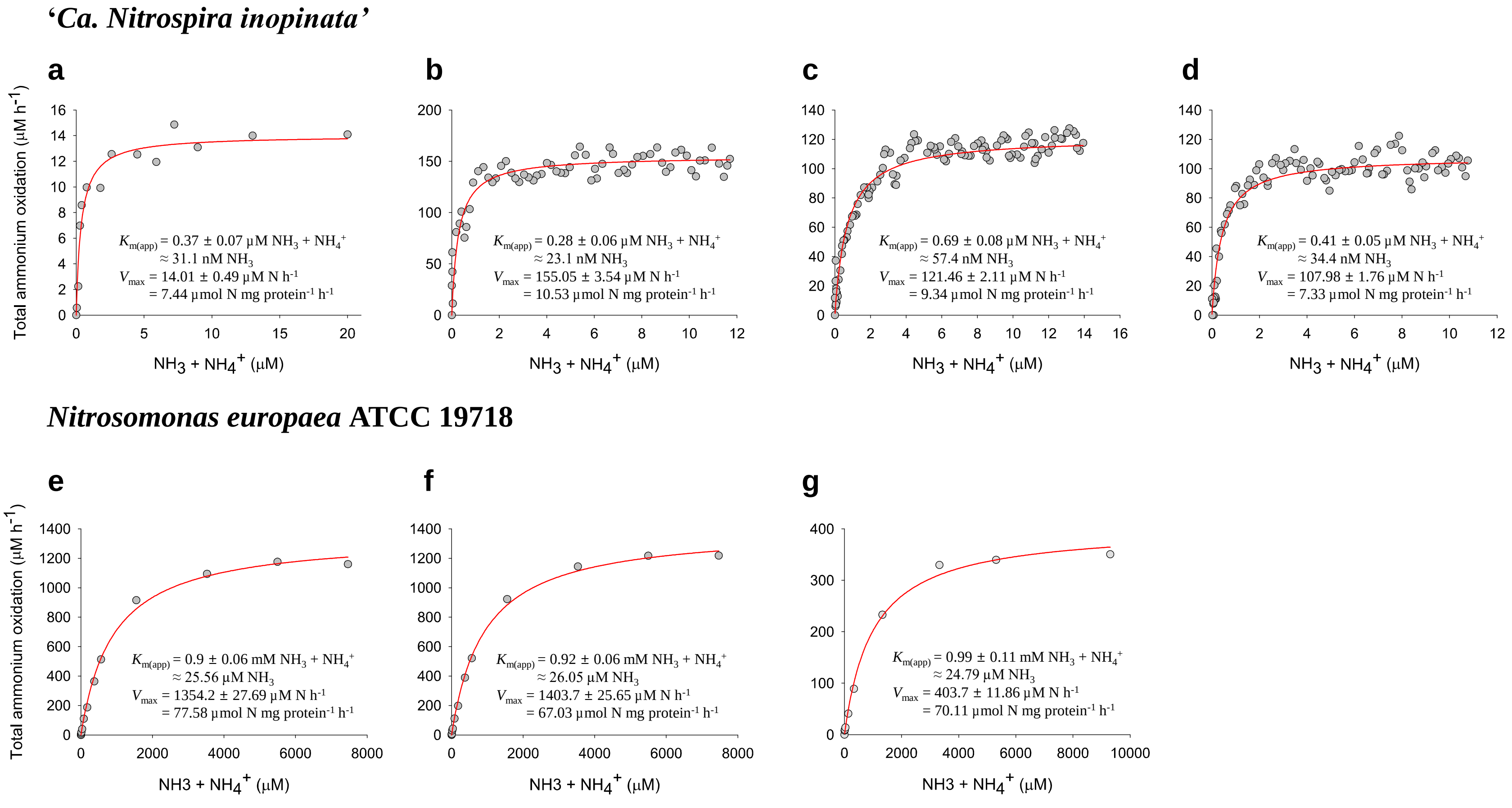
